# Integrating protein copy numbers with interaction networks to quantify stoichiometry in mammalian endocytosis

**DOI:** 10.1101/2020.10.29.361196

**Authors:** Daisy Duan, Meretta Hanson, David O. Holland, Margaret E Johnson

## Abstract

Proteins that drive processes like clathrin-mediated endocytosis (CME) are expressed at various copy numbers within a cell, from hundreds (e.g. auxilin) to millions (e.g. clathrin). Between cell types with identical genomes, copy numbers further vary significantly both in absolute and relative abundance. These variations contain essential information about each protein’s function, but how significant are these variations and how can they be quantified to infer useful functional behavior? Here, we address this by quantifying the stoichiometry of proteins involved in the CME network. We find robust trends across three cell types in proteins that are sub- vs super-stoichiometric in terms of protein function, network topology (e.g. hubs), and abundance. To perform this analysis, we first constructed the interface resolved network of 82 proteins involved in CME in mammals, plus lipid and cargo binding partners, totaling over 600 specific binding interactions. Our model solves for stoichiometric balance by optimizing each copy of a protein interface to match up to its partner interfaces, keeping the optimized copies as close as possible to observed copies. We find highly expressed, structure-forming proteins such as actin and clathrin do tend to be super-stoichiometric, or in excess of their partners, but they are not the most extreme cases. We test sensitivity of network stoichiometry to protein removal and find that hub proteins tend to be less sensitive to removal of any single partner, thus acting as buffers that compensate dosage changes. As expected, tightly coupled protein pairs (*e.g.* CAPZA2 and CAPZB) are strongly correlated. Unexpectedly, removal of functionally similar cargo adaptor proteins produces widely variable levels of disruption to the network stoichiometry. Our results predict that knockdown of the adaptor protein DAB2 will globally impact the stoichiometry of most other cargo adaptor proteins in Hela cells, with significantly less impact in fibroblast cells. This inexpensive analysis can be applied to any protein network, synthesizing disparate sources of biological data into a relatively simple and intuitive model of binding stoichiometry that can aid in dynamical modeling and experimental design.

## Introduction

The copy numbers of distinct proteins have now been counted for a number of single cells^1–5^ and tissues^6,7^, where abundances can vary per cell type by at least 6 orders of magnitude. This protein copy number data is a valuable resource for identifying function and phenotype per cell. Highly abundant proteins, such as the chaperone ATPase HSC70, often exhibit broad functionality^8^. Protein copy numbers or expression profiles indicate variations from cell-type to cell-type, or between healthy and diseased cells^9^, where the genes that encode specific proteins are otherwise identical. However, pair correlations only scratch the surface of the information available. The relative abundance of proteins reflects a network of relationships. Can we use such a network description to capture the higher order connectivity and thus quantify the significance of copy number variations on function? In clathrin-mediated endocytosis (CME), an essential process for transport across the plasma membrane^10–14^, clathrin is a highly abundant protein, but it has dozens of binding partners that could leave it in short supply. Some proteins, such as dynamin 3, are only expressed in a subset of mammalian cell types, but given that dynamin proteins are encoded by three genes, the two other genes could compensate for this shortage. Here, we quantify the relative stoichiometry of proteins in CME using a mathematical optimization that integrates both the protein network and the known copy numbers of proteins, identifying which proteins are in excess supply (super-stoichiometric), relative to their partners, and vice-versa (sub-stoichiometric). Coupled with computational ‘knockdowns’ of each protein, this approach allows us to identify strongly correlated partners and protein clusters, determine trends in stoichiometry across cell types, and predict how experimental knockdowns may impact binding interactions in CME.

Stoichiometric balance is a simple but useful way to define a quantitative model of relative protein copy numbers. For obligate multi-protein complexes in stoichiometric balance, each protein subunit is expressed at copy numbers relative to their stoichiometry in the complex. That is, if each subunit binds 1:1, they would all have identical copy numbers. The largest benefit of stoichiometric balance is an improvement of complex yield, as subunits are not sequestered in incomplete complexes^15–17^. With all interfaces matched up to a partner, perfect balance also minimizes the formation of mis-interactions that can result from leftover subunits that assemble in nonfunctional complexes due to the sticky hydrophobic surfaces of proteins^18–20^. Experimental evidence supports stoichiometric balance for proteins involved in highly stable complex formation^2,21,22^. We recently extended this idea of stoichiometric balance to larger networks where each protein may have interfaces with multiple competing partners^23^. For large reversible networks to be in balance, each interface of a protein would be expressed to match (by summing over) all of its partner interfaces. Applied to the CME network in yeast, this study found that the proteins in CME are largely in stoichiometric balance, judged relative to randomly sampled copy numbers, while specific classes of proteins, like enzymes, are sub-stoichiometric^23^. The motivation for stoichiometric balance applied to large networks is similar as for obligate complexes—one can directly quantify when specific proteins deviate from this simple underlying model that maximizes matched protein complexes. Hence this optimization problem is not a black box: each prediction of stoichiometry can be traced back to a protein’s network of partners and copy numbers to determine why observed copy numbers are sub- or super-stoichiometric relative to perfect balance.

Our model of stoichiometric balance asserts that perfect balance of each binding interface to its partners is optimal. We thus include in our interpretation of the balanced solutions the more complex hypothesis that perfect balance can be suboptimal. For example, the model effectively ignores that some binding interactions are weak and are not designed to be in permanent complex with a partner. Many of the protein-protein interactions (PPIs) that drive self-assembly, including those in CME, have affinities of only micromolar strength [see e.g.^24^ and refs within], and super-stoichiometric proteins would help drive binding and total complex yield. The model further assumes a static picture of all complexes being formed simultaneously, lacking spatial control of protein interactions^25,26^ or temporal sequences of binding events^10,25^. In the dynamic cell environment, proteins that are sub-stoichiometric can help control signal transduction and cell fate by selectively activating only one of many partners^27^. Strategic imbalances in copy numbers can also dramatically improve assembly yield^28^, by preventing sequestration of subunits in mis-interacting intermediates. In a kinetic model of clathrin vesicle formation in yeast, our recent study found that stoichiometric balance of adaptors to clathrin would produce faster vesicle formation relative to observed copy numbers^23^. However, one possible benefit for the observed imbalances is that by keeping adaptors at sub-stoichiometric levels, they can more effectively tune where and when clathrin targets the membrane, acting as gatekeepers for the speed and success of vesicle formation^23^. Hence, we will interpret sub and super-stoichiometric proteins in our network in terms of the both the costs and potential benefits provided by their mis-matched abundances.

CME involves dozens of components performing the stochastically orchestrated assembly of a protein coat for the purpose of capturing and internalizing cargo across the plasma membrane. Without an objective and quantitative framework for integrating the distinct protein copy numbers and hundreds of protein interactions, it is incredibly complicated to interpret the role of individual proteins or interactions in CME. The assumption of stoichiometric balance reduces the information required to assemble a complex network to the point that it can be reliably filled in by available experimental biochemical information. Ideally, we would model this process using physical laws that yield spatial and temporal resolution. Such dynamical modeling could predict detailed mechanisms, such as how uptake of hundreds of potential cargo proteins (e.g. G-protein coupled receptors (GPCRs) or the transferrin receptor) is controlled by the network of dozens of cytoplasmic proteins. However, this is currently intractable, given the sheer number of components and interactions, coupling to cytoskeleton^10,29–33^, membrane bending and vesicle fission^34–36^, and connection to downstream steps in vesicle trafficking. Modeling efforts have nonetheless been critical in helping quantify requirements for clathrin-cage assembly^23,37–42^, the role of the cytoskeleton^33,43,44^, the impact of dimensionality reduction on assembly^23,24^, and the mechanical coupling to the membrane^45–47^. Although the sequence of events in CME have been broadly defined^10^ (including initiation by localization of adaptors, recruitment and assembly of the clathrin lattice, and concurrent bending of the membrane), computational approaches such as ours help to predict how perturbations to single components or subsets of components will propagate throughout the network. Based on previous work^10,11,48^ the 82 proteins we define here are significant contributors to CME in mammalian cells. Moreover, they can be usefully classified, according to their established role in CME, as cargo adaptors, enzymes, structural or scaffold proteins, and cytoskeletal components. We expand our network to the membrane to include specific lipids and internalized cargo/receptors, as internalization constitutes the fundamental purpose of CME. The composition of the plasma membrane lipids impacts the membrane’s mechanical properties, but also directly controls recruitment of cytosolic proteins to the surface, particularly via negatively charged phospholipids. Further, membrane composition can be dynamically altered by enzymes that metabolize lipids, including phosphatidylinositol (PI), phosphatidylinositol-5,6-biphosphate (PI(4,5)P_2_), and later in vesicle uncoating, phosphatidylinositol-3,4-bisphosphate (PI(3,4)P₂). Sorting signals specify cargo internationalization by adaptor proteins, which couple clathrin cages to both phospholipids and cargo^49^. We show below that these functions often directly correlate with our measured stoichiometric balance, even when using copy numbers from three distinct cell types^2–5^.

A critical ingredient to stoichiometric balance analysis is that the protein interactions of the network must be resolved at the level of molecular interfaces. This level of detail is necessary to capture the ability of proteins both to compete for binding through shared domains and bind simultaneously through distinct domains. The construction of this network is laborious and although automated predictive resources exist^50,51^, they are susceptible to false positives and negatives. We assembled this interaction network for proteins involved in mammalian CME using the wealth of research detailing their domains, binding interactions, biochemistry, dynamics, and function (Table S1-S3). Our network integrates previous work annotating the domain specific binding partners of essential components such as the adaptor protein AP-2^52–54^ and clathrin^55^, with a comprehensive literature curation of 82 proteins collected from three studies of systems-level CME^10,11,56^, plus lipid and cargo binding partners^49,57^. These structurally resolved networks, on their own, provide a rich resource for studying specificity and evolution in protein interactions ^23,58–61^, and are essential for constructing dynamical models of the processes, as has been done for yeast CME^23,56^ and ErbB signaling^27^. Networks further act as maps that suggest pathways of connection and communication, highlighting, in this case, that not only proteins, but very specific interfaces mediate a majority of protein interactions. This map can therefore be useful for designing mutations or tracking how disease mutations at a specific interface might disrupt global function^51^.

Our approach here is complementary to dynamical models^42,62^, as it can be easily applied to large networks containing hundreds of interactions^23^, requires almost no parameterization (*e.g.* no biochemical data required), and provokes unique threads of questions about the relationships between proteins. The calculations are also highly efficient, taking seconds on a CPU, allowing for systematic perturbations to the network of components or their copies, across distinct cell types, with minimal expense. For example, we ask: is clathrin present in excess in the cell, like a pool available to function whenever it is needed? Are enzymes abundant and balanced in the network? Are there more adaptor proteins than available binding sites on the membrane? How do the copies of cargo add up relative to their targeting adaptor proteins? Do knockdowns (KDs) or removal of each protein result in localized or global perturbations to the stoichiometric balance and thus complex formation throughout the protein network? Rather than undergoing an extensive experimental pursuit to silence a gene encoding a CME protein *i.e.* siRNA, we show that KDs can be mimicked computationally to study their predicted effects on CME copy number distributions.

In this paper, we first introduce the mammalian proteins and their interface-resolved interaction network constructed here using protein structures, PPI databases^63^, and manual literature curation. We perform a structural and functional characterization of this network that highlights the high specificity of binding interfaces, the redundancy of domain interactions, the capacity to assemble large multi-protein complexes through multi-valent proteins, the presence of regulatory interactions, and the tight connection to the membrane via direct protein-lipid interactions. We then provide a toy example to illustrate the stoichiometric balance concept, before applying it to the copy-numbers observed in three cell types: a rat synaptosome, a mouse fibroblast, and a human epithelial (HeLa) cell. We perform this stoichiometric balance analysis to the full network, to the network without lipids and cargo, and to the network as each protein is removed or ‘knocked down’. We determine which proteins are most sensitive to knockdown of others, and how this varies by network connectivity (i.e. hub proteins). Comparisons across cell-type are powerful for identifying trends, as abundances change but the network is highly conserved. Importantly, with each perturbation to the network, we can quantify changes to the full network of protein stoichiometry, without needing to focus on the response of a specific protein or observable. We discuss the role of sub-stoichiometric proteins as gatekeepers, controlling when and where cargo uptake occurs. Additionally, chemical modifiers (like kinases), whose interactions are transient but whose changes are long-lived, may be optimal when sub-stoichiometric. We discuss the role of super-stoichiometric structural proteins as large reservoirs that are readily available when needed. Finally, we conclude with the future directions made possible by the datasets generated and analyses performed here.

## RESULTS AND DISCUSSION

### The CME proteome is well defined structurally and functionally from extensive literature

We selected the 82 proteins in our CME cytoplasmic proteome (Fig 1) using a previously curated yeast CME interactome ^65^, live-cell studies ^10,66^, and comprehensive reviews ^11^. We classified these proteins and colored them in the figures of this paper according to their primary known function in CME (Fig 1). Because the physiologic purpose of CME is to internalize transmembrane receptors or cargo across the plasma membrane, our full network includes key plasma membrane lipids and transmembrane protein targets, visible as the bottom row of nodes in Fig 2. We define nine types of transmembrane receptor/cargo, each of which is selected by (bound to) one or more cargo adaptors for inclusion into clathrin-coated vesicles (Table S1) ^49,57^. In our classification scheme, we separated the central adaptor AP-2 (mustard yellow) from the other cargo adaptors (pink), because functionally, it is the most essential adaptor, without which uptake is drastically reduced^67^, and topologically, it is a heterotetramer (encoded in 5 genes—two for the alpha adaptor, Fig 1) that acts as a hub in the network. The lipids are required for the actual localization of cytoplasmic proteins to the membrane surface, where PI(4,5)P_2_ is essential^68^. Alternative to our functional classification, CME proteins can be classified based on their arrival time at sites of CME^10,11,69^.

**Figure 1.**
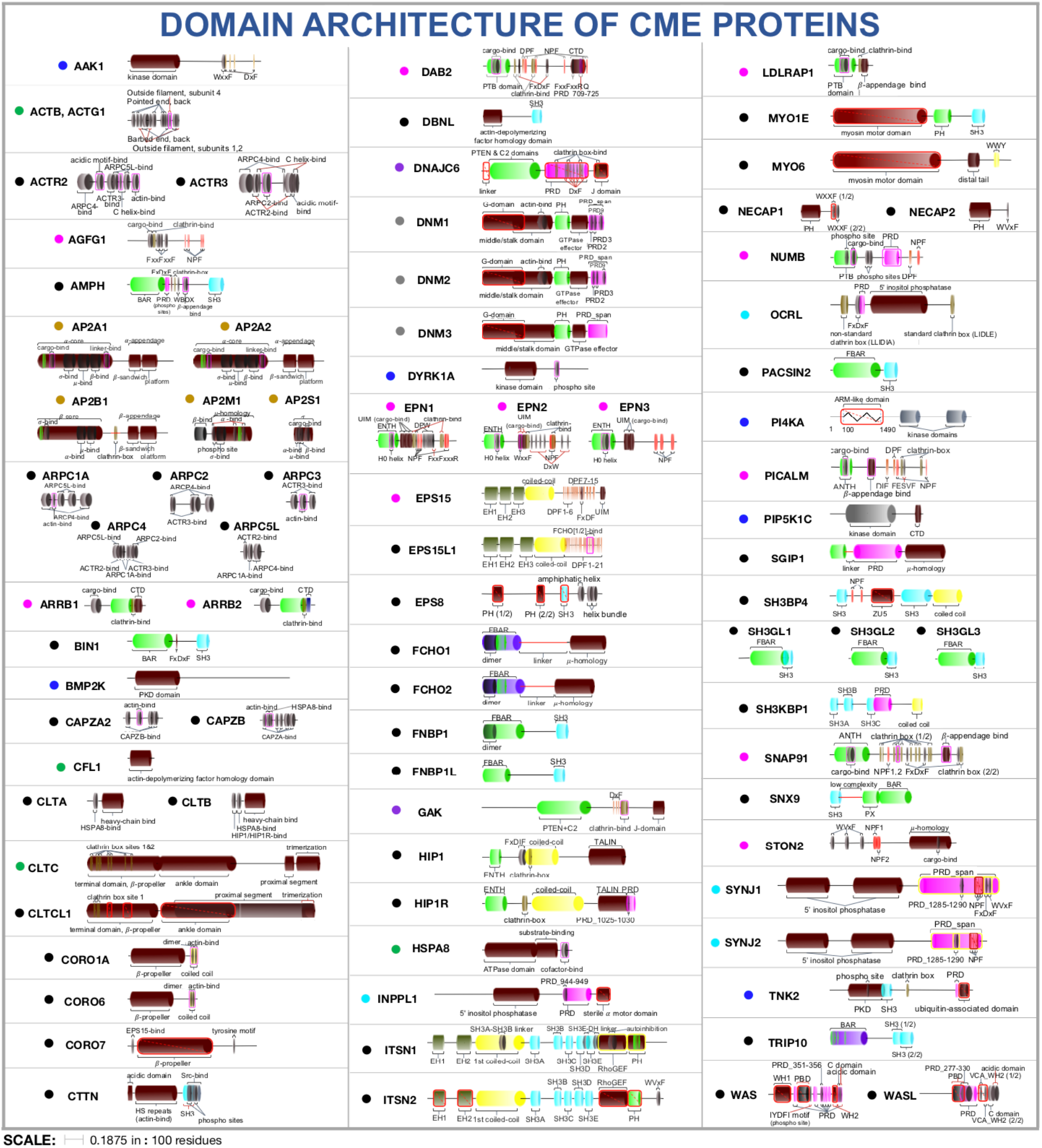
Domain architecture of 82 CME proteins in our human interface-resolved protein network. Information on domains was curated from SMART^64^ and low-throughput studies. The colored dot next to each protein name indicates the class to which we have assigned it: the central AP-2 adaptor subunits in mustard yellow, other cargo adaptor proteins in pink, kinases in dark blue (which act on proteins and lipids), phosphatases in light blue (which act only on lipids), dynamins in gray (necessary for vesicle fission), enzyme co-factors of the ATPase chaperone protein HSC70/HSPA8 in purple (needed for disassembly of cages), and a class of highly-expressed proteins in green. Protein length is scaled to reflect residue length, with distinct domains shown in colors, including structured domains and short linear motifs that mediate binding. Red outlines indicate domains that are not present in our network, as they were not assigned any interacting partners. Domains are colored: SH3 (blue), PRD (pink), lipid binding (green), coiled-coil (yellow), other structured (brown). Red boxed domains are not in our interface-resolved network due to lack of interaction partners.

**Figure 2.**
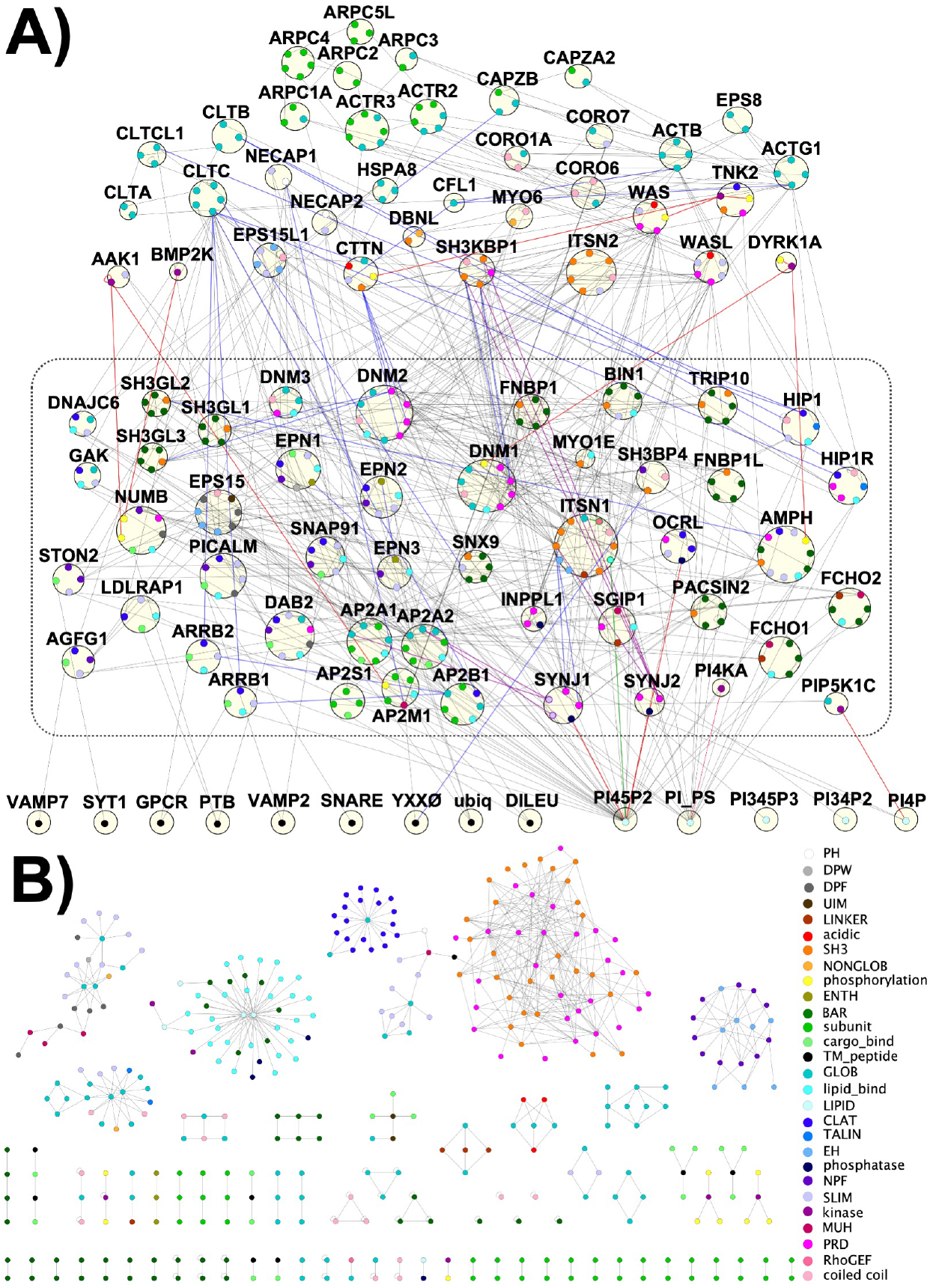
The interface-resolved PPIN for our mammalian CME proteome indicates extensive and redundant capacity to assemble and localize to the membrane. (**A**) The CME PPIN includes the 82 proteins from Fig 1, as well as 5 distinct lipids and 9 classes of transmembrane receptors or cargo along the bottom row. The box contains the set of proteins that directly bind to lipids and cargoes. Proteins that do not bind to lipids nor cargoes are located directly above the dashed black box. Unique interfaces are shown, color-labeled according to interface type. Red edges indicate enzymatic reactions, and blue edges indicate some conditionality or regulation of the interaction. Purple edges are isoform specific. (**B**) Map of interface-resolved PPIs in the network. When separated from the parent proteins, the interface-interaction network (IIN) illustrates the selectivity of binding pairs, visibly encoded in modules of pair, square, and hub motifs. **BAR**: Bin-Amphiphysin-Rvs domain; **CC**: coiled coil; **Clat.**: clathrin-box motif that mediates binding interaction with clathrin heavy chain; **DPF**: Aspartic Acid(<u>D</u>)-Proline(<u>P</u>)-Phenylalanine(<u>F</u>) motif; **DPW**: Aspartic Acid(<u>D</u>)-Proline(<u>P</u>)-Tryptophan(<u>W</u>) motif; **EH**: Eps15 Homology; **NPF** Asparagine(<u>N</u>) Proline(<u>P</u>) Phenylalanine(<u>F</u>) domain; **F-BAR**: FCH(<u>F</u>)-Bin-Amphiphysin-Rvs domain; **ENTH/ANTH**: Epsin N-terminal Homology/AP180 N-terminal Homology domain; **Glob.**: globular domain; **PH**: Pleckstrin-Homology domain; **SH3:**SRC Homology 3 domain **PRD** Proline Rich Domain; **SLIM**: short linear motif; **μHD**: mu-homology domain.

### The CME interface-resolved network can be constructed from semi-automated and manual curation

We constructed the interface-resolved network (Fig. 2) with the proteins shown in Fig 1 by downloading all documented protein-protein interactions from databases BioGrid^63^, IntAct and Mentha, and assigning interfaces to each interaction if sufficient information was available. In Methods, we detail all steps taken to construct this interface resolved network of binding interactions (see the flowchart in Fig S1), including the addition of protein-lipid (previously compiled here in ref^24^) and protein-cargo interactions^49,57,70^, as well as justification and confidence levels of our assignments. Although we added some interactions due to literature reports and homology, information was sometimes insufficient to add all possible homologous interactions. This results in some differences between homologs, for example the interactome of EPN3 and EPN1 are not identical, nor is CLTCL1 and CLTC. While this may be functionally accurate, it may also indicate a lack of independent studies or any additional support for us to make corresponding assignments. Hence, we acknowledge that our network is likely not the complete picture of the CME interactome. The proteins and copy numbers are compiled in Table S1, the binding domains in Table S2, and the full interface resolved network in Table S3.

### The interface-interaction network has a topology optimized for binding specificity

Our network has characteristics previously demonstrated to be beneficial for binding specificity, indicated directly by how the interface-interaction network (IIN) in Fig 2B breaks into islands or modules by interface type. This includes a module mediated by Src Homology 3 (SH3) and Proline-Rich Domain (PRD) interactions (orange and pink module, Fig. 2B) that we discuss further below. This modularity is expected due to constraints on evolving interfaces both for specific interactions and against nonspecific interactions^18,20,58^, which further constrains the protein network^59^. The network topology displays an abundance of hub and square motifs^59^, in addition to pairs (bottom, Fig. 2B). We defined 28 classes of interface types, some of which are structured domains (e.g. SH3, BAR), but the majority of which represent short linear interaction motifs (SLIMs), such as PRDs and NPFs. SLIMs are typically within disordered or unstructured regions of proteins, although they may adopt folds upon binding^71^ (Fig. 2B). The modules do not all separate cleanly, and the bridging nodes often have unusual binding or regulatory behavior. For instance, SH3BP4 has an SH3 domain that binds the YXXØ cargo peptide, connecting the SH3-PRD module to modules on the upper left. This upper left corner also contains a white PH domain that brings together multiple modules. The PH ear-domain of NECAP1 displays significant promiscuity in the types of partners it binds, including SLIMs, linker regions, the clathrin-box motif of AP2B1, and the AP2M1 subunit. This promiscuity allows NECAP1 to localize to sites of CME, and then act as a negative regulator of AP-2 activity^72^.

### The CME network partitions via connectivity to the membrane

By arranging our network with the membrane cargo and lipids along the bottom, we can clearly see the extent to which the cytoplasmic proteins in CME directly localize to the membrane surface, as indicated by the boxed region in the center (Fig. 2A). These proteins also contain large numbers of PPIs, allowing them to form a large variety of connected networks with multiple links to the membrane surface. The links are typically weak, which allows them to dynamically remodel throughout the formation of the clathrin coat, behaving in a manner similar to liquid droplets^73 74^. By localization to the membrane, these proteins can also exploit dimensionality reduction in what is effectively the 2D space of the surface, which stabilizes their interactions either with other proteins or to the surface, allowing proteins to nucleate complexes at much lower concentrations than are required in solution^24^. The interconnectedness and redundancy in this network is one reason why making predictions about knockdowns of CME proteins is challenging. The composition of adaptors and accessory or scaffold proteins per clathrin-coated vesicle is not unique. Apart from clathrin being essential, and AP-2 is reliably present, cargo compositions vary^75^.

The top tier of proteins, including clathrin, connect to the membrane only indirectly via protein-protein interactions. Because clathrin does not interact directly with the membrane, it uses its three n-terminal ‘feet’ to bind to one each of 14 possible distinct adaptor or accessory proteins, 13 of which also directly interact with membrane lipids (Fig. 3C excluding non-adaptor proteins OCRL, GAK and DNAJC6). The Venn diagram in Fig. 3C highlights how many proteins that bind to clathrin also bind the adaptor AP-2, membrane lipids, and cargo, numbering 10. Another large cluster of 7 proteins binds both AP-2 and lipids, whereas we found only one protein, HIP1R, which binds to clathrin and lipids but not AP-2. Thus, almost all lipid binding proteins that bind clathrin also bind AP-2, but the reverse is not true. All cargo binding proteins localize to the membrane via either direct lipid binding or via interactions with AP-2, with SH3BP4 being the only exception; SH3BP4 binds the transferrin receptor in a nonstandard way, using an SH3 domain^76^. The lipid binders that are independent of AP-2 and clathrin are almost all either lipid phosphatases or BAR-domain containing^77,78^ curvature sensors/inducers. The dynamin proteins, despite their centrality to CME^34^, do not directly bind either clathrin or AP2, as we discuss further below.

**Figure 3.**
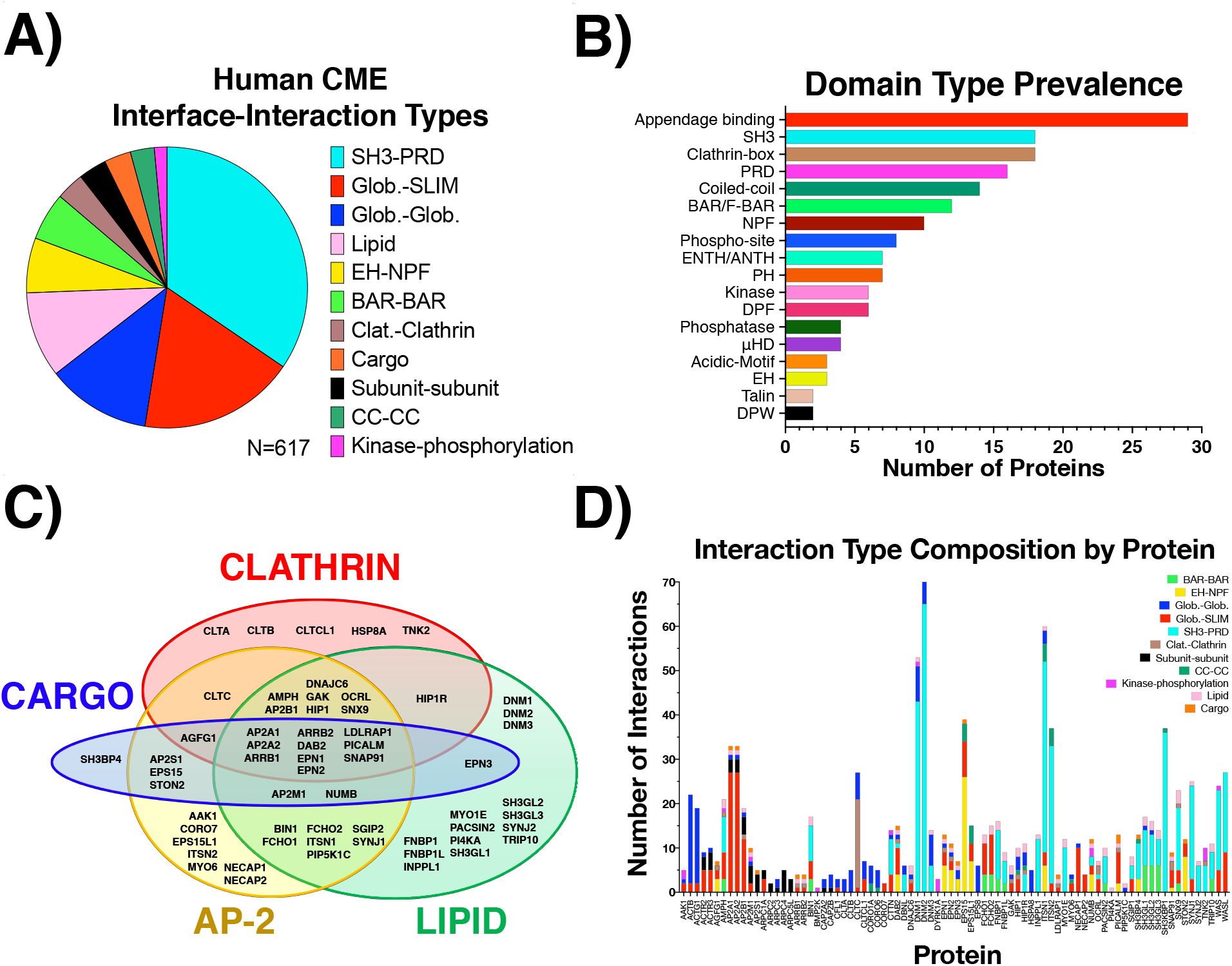
Domain types and interaction pairings in the network are dominated by binding mediated by short linear interaction motifs (SLIMs) (**A**) The types of interaction pairings in the human CME network. PRDs, SLIMs, NPF, and Clat. categories all involve SLIMs binding to structured/globular domains. (**B**) Prevalence of domain types across all human CME proteins. Counts generated if protein has at least one domain of each type present in the CME IIN. Appendage binding, PRD, clathrin-box, NPF, DPF, acidic motifs, DPW are all SLIMs, (**C**) The majority of proteins in the network bind to at least one of the functionally critical components of the AP-2 complex, clathrin triskelion, lipid, and/or cargo. Proteins that are not present in this core set are cytoskeletal. (**D**) Interface-interaction composition profile of human CME proteins, listed alphabetically along the x-axis. Counts generated if protein contains either domain involved in interface-interaction listed. Self-interactions were counted once. Domain names are defined in Fig. 2.

The remaining components of the top-tier that are not part of our defined AP-2/Clathrin/cargo core (Fig. 3C) are primarily associated with the actin cytoskeleton. In fact, of the cytoskeletal components in our network, only CORO7 and MYO6 directly interacts with AP-2 or clathrin. Instead, the cytoskeleton most directly links to the central CME machinery via the 3 dynamins. Dynamins bind directly to the essential cytoskeletal actin proteins ACTB and ACTG, which are also network hubs. The cytoskeletal helps to bend the membrane in CME, although in mammalian cells it is not required^79^.

### The majority of binding interactions are mediated by weak or transient connections involving a short linear motif

We defined 11 major interface-interaction classes based on the interface present in our network (Fig. 3). Of the 617 interactions present in the IIN, more than half involve SLIMs (Fig 3A). Of the categories we classified, PRD, Asn-Pro-Phe (NPF) motifs, and Clathrin binding motifs (listed as Clat.) are all SLIMs, as are those included in the general SLIM-Glob. (short for Globular) category such as acidic, Asp-Pro-Phe (DPF), Asp-Pro-Trp (DPW), coiled-coil (CC), linker, and ubiquitin-interacting motifs (UIM). The most common specific interaction pairs are the PRDs binding to the structured SH3 domains (Fig. 3A).

A hierarchy of connections connect clathrin with the membrane. Most CME proteins have domains that mediate interactions with AP-2, clathrin-box motifs, and SH3 (Fig. 3B). Of the 31 PRD- and/or SH3-containing proteins, 11 (AMPH, BIN1, DAB2, DNM1/2/3, HIP1R, ITSN1, MYO1E, NUMB, and SGIP1) contain lipid-binding interfaces. SH3-containing proteins mediate low-affinity hydrophobic interactions with K_*D*_s ranging from 1-10μM^80^. Interactions between the AP-2 appendage domains and the SLIMs that bind to it are also typically in the μM regime^52,53^ (see Table S2). These low affinity interactions allow for the rapid assembly and disassembly of transient complexes and clusters, as has been shown for the FCHO1-EPS15 interactions that nucleate sites of clathrin-coated pits^73,74,81^.

From the same interaction composition profile, we observe proteins that mediate much fewer interactions often conduct specific roles in phosphorylation or assembly of higher-order protein complexes (Fig. 3D). For example, ARPC2 and ARPC5L operate only as subunits of the larger ARP2/3 complex hence their PPIs are only mediated with other ARP2/3 subunits. BMP2K, PI4KA, and PIP5K1C are kinases that mediate just two, if not one, binding interactions (Fig. 2A, 3D). The behavior of kinases is distinct, performing nearly irreversible modifications that usually influence binding of subsequent proteins. Their involvement in the CME network is much less pronounced than in a signaling network^23,27^. The most significantly connected enzymes in the CME network are lipid phosphatases, which can regulate the stickiness and mechanical properties of the plasma membrane^82,83^.

### Dynamin centers a module that is differentiable from the AP-2/Clathrin/Cargo module

The interaction composition profiles for dynamin proteins, DNM1/2/3, are overrepresented by SH3-PRD. For 43 of 53 total interactions mediated by DNM1 and 65 of 74 interactions by DNM2, respectively, ^~^81% and ^~^88% of their interactions are mediated by the multiple PRD motifs within dynamin, driving their high-ranking in protein connectivity (Fig 3D). Dynamins contain no SH3 domains themselves, but use their PRDs to bind SH3 containing partners and use their GTPases activity to induce vesicle fission of clathrin-coated pits^35^. Dynamins provide a hub of a distinctive module of the CME network because they do not bind AP-2 or clathrin. Instead, they exert their influence on the CME stages via lipid-binding, acting as hub proteins to connect to several other exclusively lipid-binding proteins like SH3-containing endophilins and formin-binding proteins, as well as cytoskeletal proteins (Fig 3C). They thus play a central role in reshaping the membrane during invagination^35^.

SH3-PRD domain interactions thus act as another way to usefully partition the CME proteins by how they connect between the dynamin and AP-2/clathrin/cargo module. The SH3-PRD interactions are not used by AP-2 or clathrin, neither for cargo nor lipid binding (except SH3BP4). They cluster with lipid binding (endophilins and formin binding proteins) and cytoskeletal proteins (WAS, WASL, MYO1E, CTTN, DBNL). We can define distinct classes of proteins that crossover between the AP-2/clathrin/cargo module (Fig 3C) and the SH3-PRD module. The proteins DAB2, NUMB, SYNJ1, OCRL, SGIP1, and HIP1R all connect to the AP-2/clathrin/cargo module and have PRDs. The proteins AMPH, Bin1 (amphiphysin II), SNX9 and, SGIP1 all connect to the AP-2/clathrin/cargo module and contain BAR domains and SH3 domains. Finally, intersectins (ITSN1 and ITSN2) connect to both modules and are unique in also containing EH domains that bind to several other adaptors, providing direct links to AP-2, other adaptor proteins, dynamins, and links to the cytoskeleton.

### Measuring stoichiometry of components highlights the broad distribution of single-cell protein abundances

We quantify stoichiometry by integrating the interface-resolved protein network with copy numbers derived from single cell types^2,3^. The model we use defines stoichiometrically balanced copy numbers as matching up each copy of an interface to one of its partners^23^. The model thus solves for an optimal equilibrium distribution of bound complexes in the limit that all binding interactions are equally strong. We constrain the balanced copy number solutions to be as close as possible to the observed, real copies for a protein. For multi-interface proteins, we further constrain each domain within the proteins to have similar copy numbers (*i.e.* a specific protein should not have 1 million copies of one domain, and only 10 of another); we allow variance due to known fluctuations in protein copy numbers^1^. The model is solved using nonlinear optimization performed by the SBOPN method (Stoichiometric Balance Optimized for Protein Networks), which takes <1 second (see Methods) ^23^. To illustrate, we show a simple IIN in Fig. 4A with observed protein copies listed. We then apply the SBOPN method, resulting in a solution of balanced copy numbers (Fig. 4B). We define the stoichiometric balance ratio, or SB ratio, of a protein *p* as the ratio between observed (*C*_obs_) and balanced (*C*_balanced_) copy numbers, which determines whether a protein is sub- or super-stoichiometric.

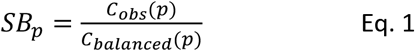

Perfectly stoichiometric proteins thus have an SB ratio of 1. Proteins that have too few observed copies, or are in demand, are sub-stoichiometric (SB ratio<1) Proteins with too many observed copies, or have excess supply, are super-stoichiometric (SB ratio>1).

**Figure 4.**
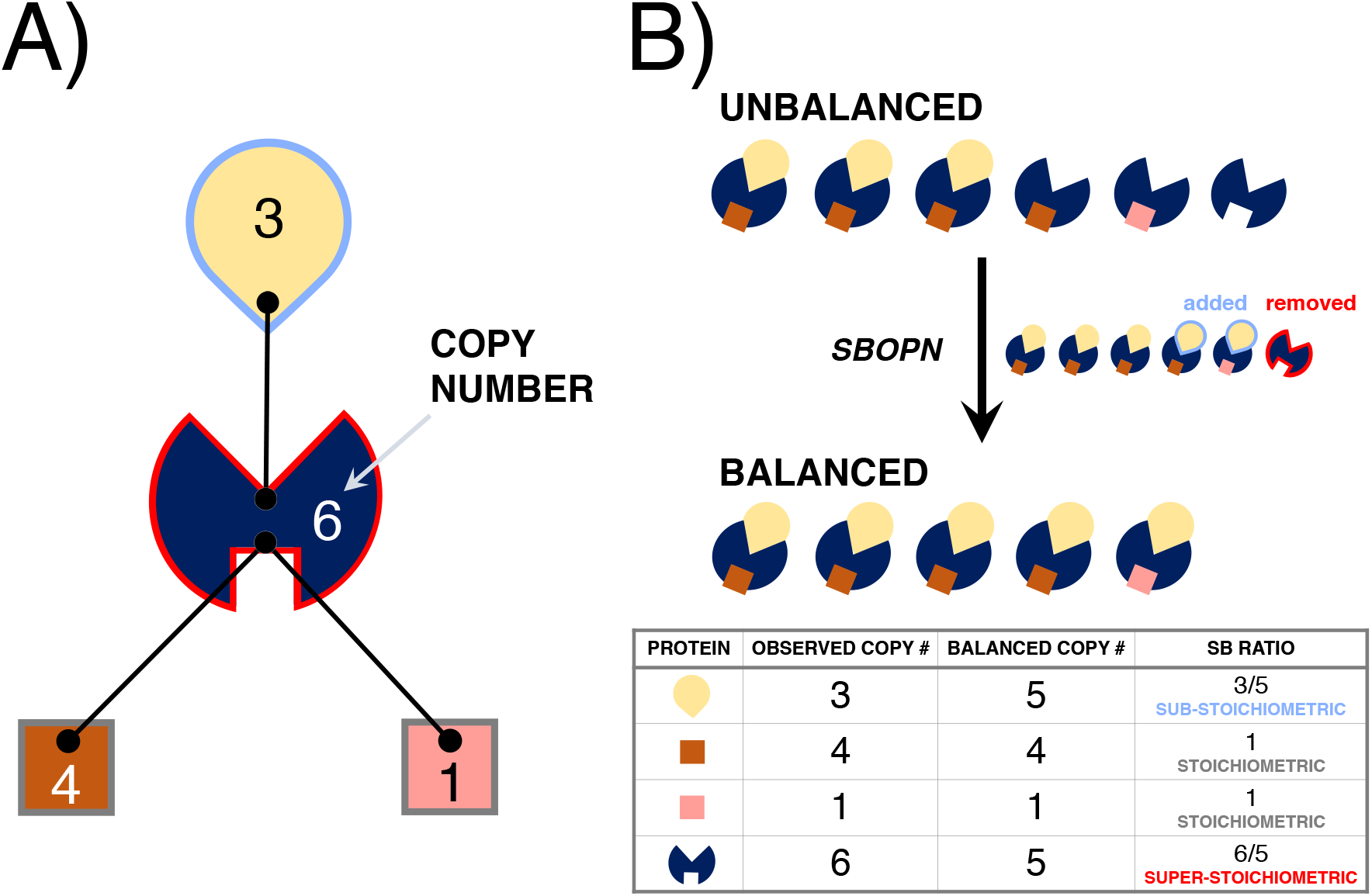
Illustration of the Stoichiometric Balance Optimization of Protein Networks (SBOPN) model applied to an interface-resolved network with observed copy numbers. (**A**) A simplified network diagram that contains four proteins, differentiated by color, where the central protein has two interfaces, and is thus capable of multi-component assembly. In the network, interfaces represented by black dots can only bind to their partners if their shapes are compatible, as reflected in nature where protein binding involves conformational and chemical compatibility. Edges represented in black lines denote binding interactions. Observed copy numbers for each protein are shown in white/black text. (**B**) The observed copy numbers are unbalanced: several interfaces are unmatched. Note here that the square proteins compete for the same binding interface. Stoichiometric balance (SB) is achieved if all interface copies are matched to their partners and there are no leftovers. Here, the SBOPN algorithm removes an extra copy of the hub protein and adds two additional copies of the droplet-shaped protein to completely match all interfaces. To quantify SB, we calculate SB ratio = observed copies/balanced copies, as illustrate in the last column of the table (Eq 1). We note that for multi-interface proteins (*i.e*. the blue one), not all interfaces must appear at the same copy number. When calculating SB ratio for a protein, we thus use the copies averaged across all interfaces.

The observed copy numbers for the HeLa cell type, shown in Fig. 5A, span multiple decades, from high pM for dynamin1 to >10μM for cofilin, with the central lipids PI(4,5)P2 and Phosphoinositide/Phosphoserines reaching even higher concentrations. Although lipids and cargo are restricted to the 2D membrane, we can define their concentration via copies/cell volume to enable interface matching in the SB analysis. These copy numbers are averaged across two studies, with high correlation (R=0.83) between the studies (Fig. S2). We note there is a distinction in whether a protein has zero observed copies, or whether the number of copies is unknown. Proteins with unknown copy numbers remain in the network and are subject to stoichiometric balancing, but without any constraint on their target copy numbers. Those proteins not observed in a given cell type (according to the Human Protein Atlas and the Proteomics DataBase) are removed from the network for that cell type, given that they are not expressed (see Methods, Table S1). Importantly, the optimal SB for any protein in a network depends on the global network and copy numbers, and thus will vary by cell type and with addition or deletion of nodes to the network. We provide all code and input files for this analysis at github.com/mjohn218/StoichiometricBalance, including code for systematically simulating knockdowns.

**Figure 5:**
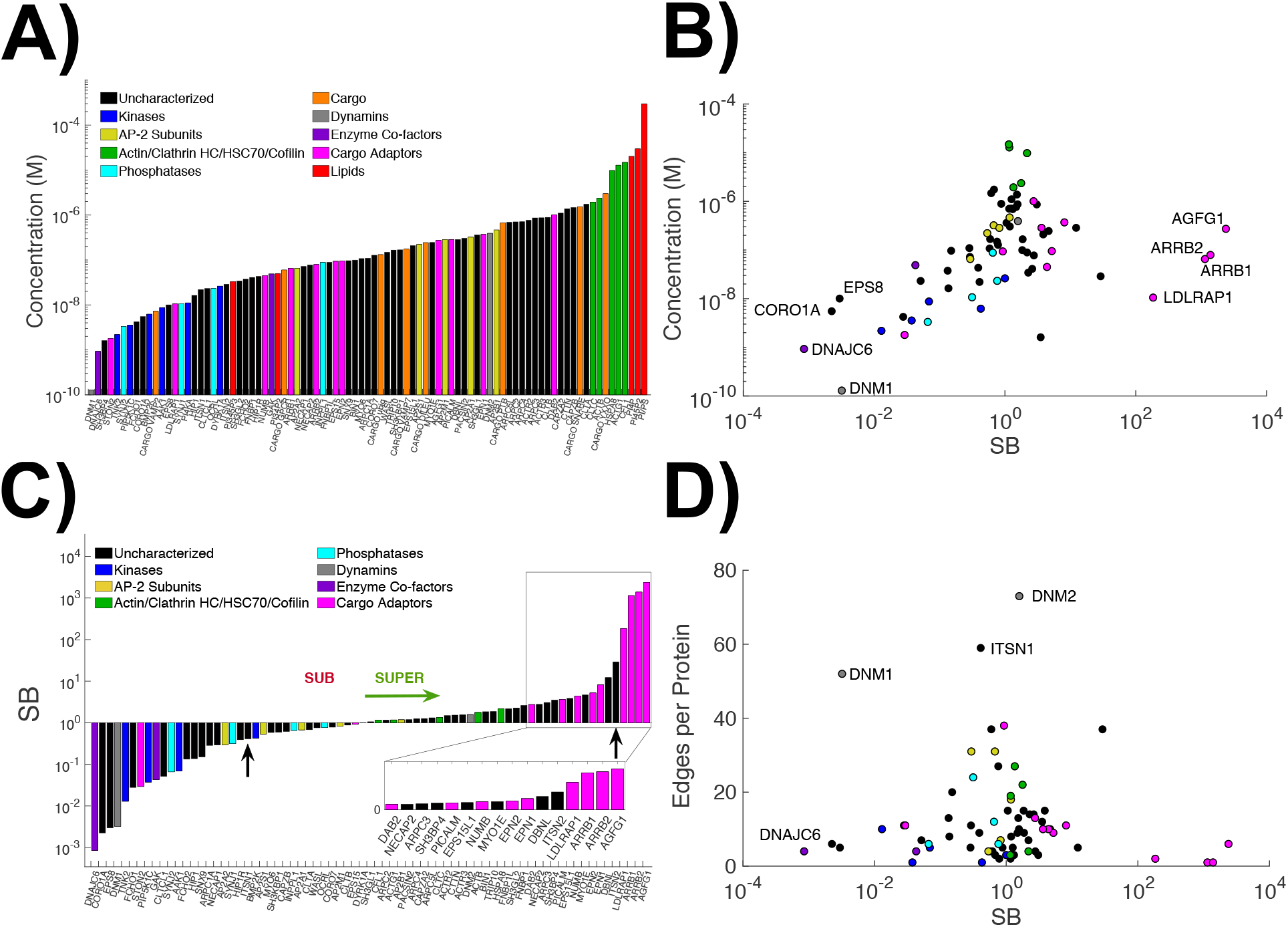
CME protein copy numbers in HeLa cells span decades, but balanced copies show functional and topological trends. **A)** Proteins, lipids, and transmembrane cargo are colored according to our classification scheme shown in the legend, and sorted here along the x-axis by copy number. The green class was designed to capture the most highly expressed cytosolic proteins, shown as a cluster on the right-hand side. EPN3 is included in the SB analysis, although its copy numbers are unknown. Seven proteins were removed from the network for having zero expected copies in HeLa (AMPH, CORO6, DNM3, SGIP1, SNAP91, WAS, CARGO_SYT1). **B)** After applying SBOPN, the SB ratios do show correlation to the observed copy numbers: proteins left of 1on the x-axis are sub-stoichiometric and tend to have low concentrations (R=0.45). Highly expressed proteins are more likely to be stoichiometric, or super-stoichiometric (green cluster). **C)** By sorting the proteins according to increasing SB ratio, we see that the cargo adaptor proteins are nearly all super-stoichiometric, or expressed at higher levels than they are ‘needed’. **D)** The SB ratios also show correlation with the connectivity or edge numbers per protein, but in a volcano shape. Highly connected proteins tend to be stoichiometric, whereas proteins with fewer connections span the full range of SB ratios.

### Stoichiometric Balance ratio correlates with abundance and network connectivity

We find that the correlation of high protein abundance with super-stoichiometry is significant, with R=0.45 (Fig 5B). While this is perhaps not surprising, our network analysis also identifies highly expressed proteins that are *not* super-stoichiometric, due to, for example, their large number of interaction partners. We find that the strongest predictor is that proteins with low expression levels are sub-stoichiometric. This is true of all the kinases and phosphatases in the network, all of which exhibit concentrations <100nM. The concentration of kinases is especially low. The co-factors auxilin (DNAJC6) and GAK, of the chaperone HSC70 (*HSPA8*), are both sub-stoichiometric, particularly auxilin which is present at only a few hundred copies. Their highly mismatched stoichiometry is exacerbated by the fact that they both bind to the highly expressed proteins HSC70 and clathrin heavy chain (CLTC). This same trend with kinases/phosphatases being sub-stoichiometric was observed in yeast ^23^. Because transient enzyme interactions result in long-lived chemical modifications to substrates, a one-to-one stoichiometry of enzymes with targets would seem unnecessary. The set of highly expressed proteins labeled in green in Fig 5A do tend to be super-stoichiometric, although they are not the most extreme (Fig 5C). Algorithmically, it is more costly to create a significant imbalance at these high abundances, and thus they are closer to an SB ratio of 1. Mechanistically in the cell, this correlates nicely with the essentiality of these specific proteins (clathrin, actin, and HSC70); although they are highly abundant and super-stoichiometric, they are not in excessive surplus relative to their full network of partners, but are closer to the ‘right’ balance.

The trend between SB ratios and connectivity in the network is visible as a volcano shape (Fig. 5D), with more highly connected proteins having an SB ratio closer to 1. Mechanistically, highly connected or hub proteins are more likely to be essential for cell survival^84^, and having an SB ratio close to 1 suggests that the proteins are present at the optimal abundance to interact with their many partners. However, we note that this correlation is weaker than the correlation of SB ratio with concentration. This is likely because protein abundances span orders of magnitude, having a wide impact on the overall SB distribution, whereas protein connectivity only spans two orders of magnitude (1-100). Abundance also does not correlate with connectivity, resulting in conflicting pressures on the stoichiometry of some proteins (HeLa: R=-0.08, Fibro: R=-0.02, Synaptosome: R=0.27). Dynamin1 (DNM1), for example, is highly connected but also has very low abundance. We find it to be highly sub-stoichiometric in the network, driven by its extremely low copy numbers; its high connectivity would favor balanced stoichiometry.

### Cargo adaptor proteins (aside from AP-2) compete to bind the AP-2 and clathrin hub interfaces

When we apply SBOPN to the CME network restricted to the 82 cytosolic proteins, we observe that nearly all of the non-AP-2 cargo adaptor proteins are classified as super-stoichiometric, that is, present at levels exceeding their partners’ demand (pink bars in Fig. 5C). Yet they exhibit copy numbers across the spectrum of concentrations (Fig. 5A). Instead, it is their network connectivity that determines super-stoichiometry. Connectivity is similar across all cargo adaptors and dominated by links to both the AP-2 appendage hubs and the clathrin N-terminal hub. There is too much competition for adaptor proteins to bind to the hub interfaces on AP-2 and clathrin, and insufficient hub protein interfaces to fulfill stoichiometric balance. Thus, we find that although clathrin is abundant, its many partners compete for binding at its N-terminal hub. Because these partners use short motifs to bind clathrin^85^, they are expected to have similar affinities. Cargo adaptor protein abundance, therefore, should play a strong role in controlling their stoichiometry with clathrin. In order to balance out the interfaces to one another, our analysis indicates the AP-2 subunits are insufficiently available, resulting in sub-stoichiometry. Nearly all the other adaptor proteins, in contrast, are in excess relative to their observed copies, resulting in super-stoichiometry. For the adaptors ARRB1, ARRB2, LDLRAP, and AGFG1, they are set to the minimum possible copies, or effectively ‘sacrificed’ in the optimization; this is due to their similar binding profiles in the network, as unlike DAB2 and PICALM, they contain almost no links outside of clathrin, AP-2, cargo, and lipids. We find exceptions to the super-stoichiometry of cargo adaptors with STON2 and EPS15, both of which are sub-stoichiometric, and neither of which binds clathrin. Their distinct network connectivity to AP-2 but not clathrin differentiates them from the other adaptors. As we show below, by removing nodes or ‘knockdowns’, one can more clearly see correlations between proteins and differences between members of each class.

### Homologs do not always display the same stoichiometry

The homologs FCHO1 and FCHO2, while not identical in their binding profiles, show similar sub-stoichiometry in HeLa cells (Fig 5C). There are, however, homologs that display differential stoichiometric ratios. For dynamin1 and 2, this is attributable to their vast differences, by a factor of 3000, in copy numbers. Intersectin 1 (ITSN1) and 2 (ITSN2) have very different SB ratios (black arrows, Fig. 5C), yet have only minor differences in their moderate-scale copy numbers, with ITSN1 at 22 nM and ITSN2 at 29 nM. It is further surprising to see the observed differences given the similar (and diverse) domain architecture of these highly connected hub proteins (Fig. 1). However, the domains exhibit different binding selectivity. We found that ITSN1 uses 12 distinct binding interfaces, totaling 62 binding interactions, whereas for ITSN2, only 7 distinct interfaces maintain 38 binding interactions (Fig. 2). They both have similar binding profiles for their SH3 domains. Hence, the main driver of the different stoichiometries are the EH (Eps15-Homology) domains, which ITSN1 uses to bind multiple, highly expressed adaptor proteins, including DAB2. ITSN1 is thus sub-stoichiometric. ITSN2, on the other hand, has not been shown to bind to the SLIM motifs in the CME adaptor proteins, despite containing two EH domains. Therefore, our analysis indicates the danger of relying on broad homology to explain the sensitivity of individual proteins to perturbations of concentration, and instead highlights the importance of individual binding domains.

### The stoichiometric balance of transmembrane cargo proteins is controlled by their cytosolic partners

Generally, SB ratios do not change significantly when the SBOPN analysis is extended to include transmembrane cargo (compare Fig. 5C and Fig. 6A). Somewhat unexpectedly, given the addition of binding partners, most adaptor proteins remain super-stoichiometric (Fig 5A). Our analysis finds that transmembrane cargo have lower abundance than their sorting adaptor proteins (Table S4), meaning there are more than enough cytosolic adaptors to select them for uptake. With no additional network connections, the transmembrane cargo thus have stoichiometry determined through their adaptors. The only transmembrane cargo we find in excess of their sorting adaptors are the abundant cargo YXXØ (which include the transferrin receptor) and SNARE cargo. They are both super-stoichiometric. Interestingly, the least abundant cargo in HeLa, VAMP2, is also super-stoichiometric, despite having only 7 nM concentration. This can be traced to the adaptor protein PICALM, the only one to select both VAMP2 and the SNARE for uptake in HeLa cells. Both these cargo are necessary for downstream vesicle fusion, and because the SNARE class is abundant at 1.5 μM, the supply of PICALM at 0.28 μM is insufficient to match up with the sum of both cargo. Thus, both its VAMP2 and SNARE cargo are in excess supply. Due to its links to both the AP-2 and clathrin hubs, PICALM is itself also super-stoichiometric. This reinforces that the SB ratio of a protein is due to its global connectivity in the network, not just its immediate binding partners. If PICALM only bound to its SNARE cargo, its observed copies would be insufficient, and we would classify it as sub-stoichiometric. The YXXØ cargo is also quite abundant at 3 μM, and its partners AP2M1 and SH3BP4 are insufficient to match its interfaces.

**Figure 6.**
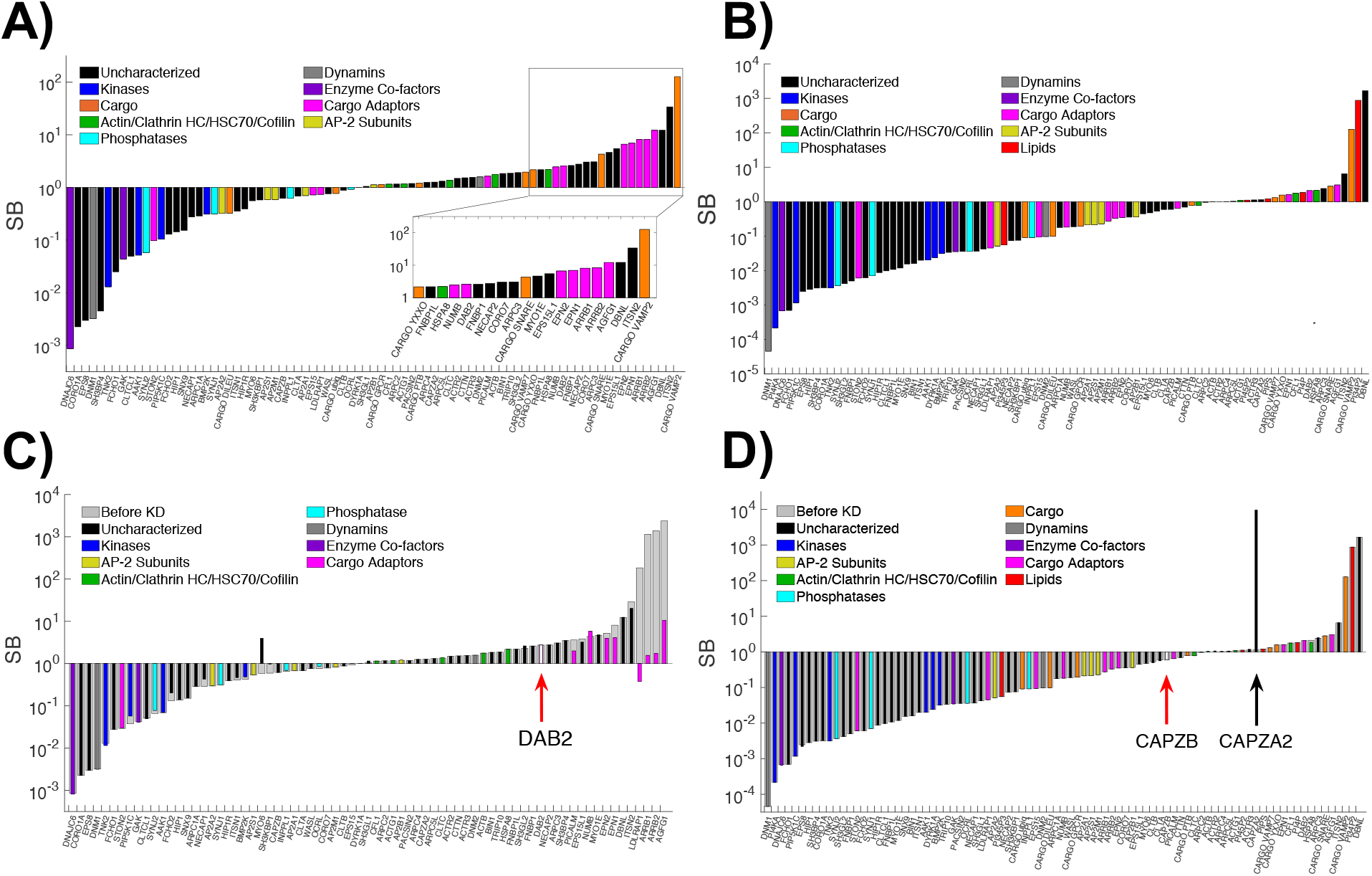
Stoichiometric balance reveals correlations and couplings in the network by adding and subtracting nodes. (**A**) Stoichiometric balance with transmembrane cargo added to cystosolic protein network. Cargo adaptor proteins are still super-stoichiometric. (**B**) Stoichiometric balance in network with all lipids and cargo added. Lipids are abundant and shift most proteins to sub-stoichiometry. **C**) Analysis of SB ratios upon removal of DAB2 protein. Light gray bars are the original SB distribution. Overlaid colored bars are SB ratios after removal of DAB2. (**D**) Analysis of SB ratios upon removal of CAPZB protein. Light gray bars are the original SB distribution. Overlaid colored bars are SB ratios after removal of DAB2.

That this cargo for the central AP-2 adaptor is in surplus suggests another mechanism for AP-2 to more stably nucleate clathrin coated pits on the membrane via its more abundant cargo. Recent experiments indicate that cargo selection is most impactful on coated pit maturation rather than initiation^69^, and establishing the timescales in future work by combining the network and abundances with spatial modeling^42^ would further clarify how interactions with cargo can checkpoint the assembly of clathrin-coated vesicles^86^.

### Membrane lipids are abundant and shift most proteins to sub-stoichiometry

When we quantify stoichiometric balance in the network with now both transmembrane cargo and lipids added in, we observe that most cytoplasmic CME proteins shift to sub-stoichiometry (Fig 6B, Fig S3) relative to before (Fig. 5A). This is due almost entirely to the hub lipids PI(4,5)P_2_ and our PI/PS class, which have very high 30μM and 300μM concentrations, respectively. It is an important caveat to note that these lipids have many other binding partners in the cell, that would thus impact their available concentrations to bind the CME proteome. However, the analysis with lipids is nonetheless instructive because not all CME proteins do bind the membrane (Fig 2A). Further, we can increase the binding stoichiometry of proteins:lipids in our analysis from 1:1 to 1:10, for example (Fig S3) which reduces the impact of lipids on shifting the network stoichiometry. We find, as expected, the upper tier of proteins (Fig 2A) which are separated from the membrane are less impacted by the addition of lipids. Instead, the most dramatic changes occur in the cargo adaptor proteins, and the BAR domain containing proteins that bind lipids and dynamin (Fig 3C). The cargo adaptors that bind lipids are now sub-stoichiometric, unlike before (LDLRAP, NUMB, ARRB1, ARRB2, PICALM, EPN2), with AGFG1/Hrb being the only exception because, rather unusually, it does not bind lipids, only its cargo. The BAR domain proteins (SH3GL2, FNBP, BIN1) all bind to the PI/PS lipid class, and switch to sub-stoichiometric upon addition of lipids, whereas the BAR proteins that bind to AP-2 (FCho1/2) are less affected, as they were already sub-stoichiometric due to low copy numbers and connections to AP-2. For much of the analysis below, we remove the lipids from the network to query selective interactions between proteins, as the high abundance of the highly connected lipids tends to attenuate relationships between cytosolic binding partners.

### DAB2 knockdown induces a global effect on CME balanced copy number distribution

By computationally removing each protein from the network, we can clearly see how a single protein virtual knockdown can cause pronounced global or local effects on the SB ratios of the full protein network, or have no detectable impact. The most striking result we find is the response to DAB2 KD (Fig. 6C) which globally shifts the stoichiometry of the other adaptor proteins. DAB2 is the most highly expressed cargo adaptor (^~^1 μM). There is excess DAB2 to select its cargo PTB^87^ (0.67 μM), which is also internalized by the adaptors NUMB and LDLRAP1. DAB2 contains diverse connections to not only clathrin, AP-2, lipids, and cargo (Fig. 3C), but also EPS15, FCHO2, MYO6, ITSN1, NECAP1, and SH3KBP1 (Fig 2). We see that DAB2 KD causes Myosin 6 (MYO6) to switch from being sub- to super-stoichiometric, and LDLRAP1 from super- to sub-stoichiometric. Likewise, most cargo adaptors are strongly impacted. For example, the SB ratios of ARRB1 and ARRB2 are significantly reduced (Fig. 6C). The reason is that the removal of the highly-expressed DAB2 makes available spots on the hub interfaces of AP-2 and clathrin, which can now accommodate more of the other adaptor proteins. Our computational KD thus would predict that removal of DAB2 would significantly reduce uptake of the LDL receptor (because LDLRAP1/ARH and NUMB also select it), and that uptake of alternate cargo would increase. This is consistent with experimental results that used RNAi-mediated gene silencing to knockdown DAB2^75^. They found a build-up of LDL receptor on the membrane (indicating reduced internalization), while microscopy images showed a reduction of transferrin receptor on the membrane. This reduction of the alternate cargo on the membrane would indicate increased internalization with DAB2 removed^75^. We note that this compensation in which cargo is taken up is not found with all experimental knockdowns of adaptor proteins^88^, because the adaptors also can contribute to efficient clathrin coat assembly. Removal of PICALM, for example, reduced transferrin receptor uptake^88^. DAB2 perhaps operates more independently of AP-2 to coordinate uptake of its own cargo, as it does not need AP-2 binding to internalize LDL receptors^75^.

### Capzb knockdown induces a localized effect on balanced copy number distribution

Unlike the global SB shift accompanying DAB2 KD, a strong local coupling emerges for F-actin capping protein subunit b (CAPZB) and F-actin capping protein subunit a (CAPZA2), which have similar copy numbers to one another (Fig. 6D). Both proteins have low connectivity in the network (Fig. 2) but contain interfaces to the highly abundant and connected actin interfaces, and CAPZB also binds the highly abundant HSC70/*HSPA8*. After applying SBOPN, CAPZA2 and CAPZB copy numbers are quite irregular from one interface to the next. As a result, when CAPZB is removed, the SB ratio of CAPZA2 changes dramatically. We find that the knockdown of CAPZB has an extreme local effect only on its immediate partner, CAPZA2. This trend is preserved in just the protein network (Fig 5A), with the addition of cargo (Fig 6AB), and with the further addition of lipids (Fig. 6D).

### Knockdown of hub proteins and highly abundant proteins is more disruptive to the network stoichiometry

By systematically knocking down each protein (or group of proteins) from the network, we can rank which proteins have the biggest impact on the SB distribution and thus complex formation in the network (Fig. S4, S5). These measurements can provide guidance in determining which proteins could be removed from the network in future modeling efforts, for example, as we would expect their influence on system dynamics to be weaker. We quantify the change upon knockdown by the distance between the distributions before and after, using the Jensen-Shannon distance, or JSD (Fig. S6 and Methods), with the most disruptive single knockdowns shown in Fig. 6C (JSD=0.7), 6D (JSD=0.55). When we sort the proteins that have the biggest impact (Fig S7), we find that these disruptor proteins have higher abundance and/or connectivity. In other words, the JSD has a positive correlation of 0.27 with protein abundance, and a positive correlation of 0.34 with protein connectivity (Fig. S8). The correlation is significant but not strong, in part because as we noted above, connectivity and copy numbers are not strongly correlated (e.g. DNM1 is highly connected but has very few copies). Also, there are a number of coupled nodes that need not have high copies or connectivity to strongly disrupt the SB ratio of one protein, with CAPZB and CAPZA2 being one pair (Fig 6d), and two components of the ARP2/3 complex (ACTR3 and ARPC3) being another (Fig. S7).

We were surprised that KD of the hub proteins of AP-2 and clathrin did not have a more dramatic impact on the SB distribution, finding a moderate impact that is maintained when all subunits of the heterotetramer AP-2 are removed (JSD=0.04), or all clathrin chains (JSD=0.05) are removed (Fig S9). While their removal did have an impact on SB ratios of many proteins (Fig S9), consistent with their broad connectivity, the JSD between the distributions was not as large as results from localized change from CAPZB, for example. We see that even with the removal of AP-2, for example, cargo adaptor proteins still have to compete with one another to bind the clathrin hub, and thus while the number of bound complexes changes, most cargo adaptors are still super-stoichiometric. The same is true of clathrin removal—many of its partners still compete to bind the AP-2 hub. Thus, when we remove both clathrin and AP-2, we see a dramatic change in the SB distribution (JSD=0.55) with several adaptors losing connections to the network entirely. While these computational KD outcomes cannot predict that removal of clathrin should stop endocytosis entirely, this is because the SB analysis reports on which complexes will be formed. Even in the absence of clathrin, the network of CME proteins would still interact with one another to form complexes, despite the fact that functional vesicles could not be produced.

### Proteins that are sensitive to the KD of others have low network connectivity

We can also quantify which proteins have unstable SB ratios, or SB ratios that are sensitive to computational KD of other proteins by measuring variance sSB,KD across all KDs (Fig. S10 and Eq 3). We find an anti-correlation between the unstable proteins, and the disruptor proteins that alter the SB distribution when they are removed (R=-0.51 Fig. S11). For example, when the disruptor protein DAB2 is removed, it induces a broad change in the stoichiometry of other proteins. However, no matter which other proteins are removed from the network, the SB ratio of DAB2 changes very little, indicating that it is robust and stable against perturbations to the network. On the other end is LDLRAP1, whose SB ratio is unstable in response to KD of an array of other proteins in the network, but whose own KD hardly disrupts the SB distribution at all. This result further highlights how despite their shared function as cargo adaptor proteins, DAB2 and LDLRAP1 have dramatically distinct impact on network stoichiometry.

The unstable proteins that are sensitive to the removal of others tend to have fewer copies (Fig. S11, R=-0.29), and have fewer connections (Fig. S11, R=-0.23). Algorithmically, changing their SB values is one of the least costly and most optimal way to balance the network stoichiometry. Mechanistically, this suggests that these types of proteins are controlled by their more highly connected partners, adapting themselves rather than inducing changes in others. Generally, unstable proteins have minimal connections in the network (Fig. S12), an observation that holds whether or not the membrane is included in the analysis. As in the previous section, the correlation is only moderate. We find that just because a protein has minimal connections in the network does not mean that it will be unstable upon KD of others. Two of the ARP2/3 subunits, for example, have only a few connections (all to other subunits), yet their SB ratios are both quite stable. This is because their partners have similar copy numbers, since they are part of a multi-subunit complex (Table S1), and these subunits do not connect to any other proteins in the network. Another subunit, ARPC3, in contrast, is also minimally connected, but it is unstable because it links outside of the ARP2/3 complex to hub interfaces on actin.

### Both protein abundances and SB ratios are similar when comparing fibroblast and HeLa cells

We can apply the same analysis to new cell types to quantify how copy number distributions and stoichiometry change in distinct cellular environments. For a fibroblast cell, the network is quite similar to that of HeLa, except one protein is added (AMPH), and five are removed due to zero observed copies (ARRB2, CTLCL1, DNAJC6, EPN3, SH3GL2). The observed copy numbers are strongly correlated between the cell types (Fig. S13) with R=0.77, although we note that the fibroblast has far more—18 total— unknown abundances. After applying SBOPN, we find that the SB ratios in the fibroblast also correlate with the SB ratios in the HeLa cells. For just the protein network, we find R=0.38, but with the addition of cargo, the correlation increases to 0.68 (Fig 7A), and to R=0.76 when the full membrane components are included. We see quite similar SB ratios for both adaptor proteins and cargo proteins across both cell types, where again we find that most adaptor proteins are super-stoichiometric (compare Fig 7A and Fig 6A). Only EPN1 has switched from super- to sub-stoichiometric in fibroblast. Although EPN1 is over 30x less concentrated in fibroblast, so is its ubiquitylated cargo, showing that our analysis reveals a correlated decrease in both EPN1 and its cargo. Its sub-stoichiometry is instead driven by binding to the AP-2 beta subunit, where more copies of it are needed, in part because DAB2 is present at much lower levels in fibroblast (14x) compared to HeLa. Hence DAB2 is much less disruptive to stoichiometry in fibroblast due to its markedly reduced copies.

**Figure 7.**
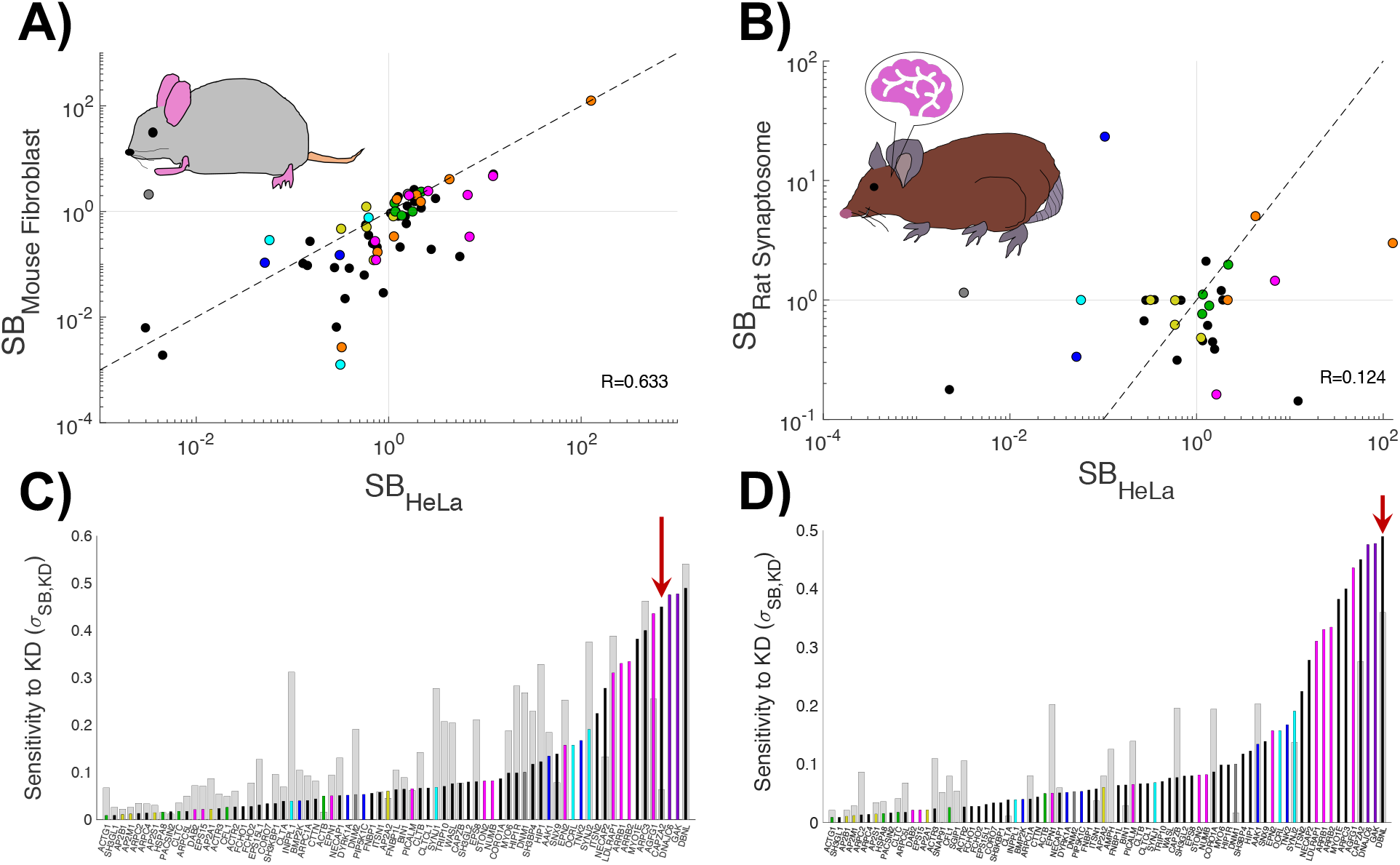
Comparison of SB values between HeLa, mouse fibroblast, and rat synaptosome cells reveals distinct patterns. (**A**) Correlation of SB ratios of proteins expressed in both HeLa and mouse fibroblast cells, has a R=0.76. The cytosolic volume of a mouse fibroblast (1200 mm^3^) was used to calculate protein concentrations (M). Dashed line shown corresponds to x=y. (**B**) Correlation of concentrations of proteins expressed in both HeLa and rat synaptosome cells. The cytosolic volume of a synaptic bouton (without vesicles and mitochondria, 0.235 mm^3^) was used to calculate protein concentrations (M). Dashed line as in (A). (**C**) Susceptibility of proteins to KD in HeLa cells versus mouse fibroblasts. Shown are standard deviations of SB ratios upon KD of other proteins in HeLa and mouse fibroblast cells. Overlaid colored bars are sensitivity measurements for HeLa proteins. Light gray bars in background are sensitivities of proteins to KD in mouse fibroblasts. Gaps indicate mouse proteins with either unknown or zero copies. Red arrow points to CAPZA2, which is not sensitive in fibroblast. (**D**) Susceptibility of proteins to KD in HeLa cells versus rat synaptosomes. Shown are standard deviations of SB ratios upon KD of other proteins in HeLa and rat synaptosome cells. Due to the predominantly low copy numbers of proteins expressed in the rat synaptosome, a linear min-max normalization was applied to their sensitivity values relative to those in the HeLa cell background. Overlaid colored bars are sensitivity measurements for HeLa proteins. Light gray bars in background are sensitivities of proteins to KD in rat synaptosomes. Red arrow points to DBNL, which is sensitive to KD in all 3 cell types. Colors as in Fig. 6A.

Despite the positive correlation between the stoichiometries and sensitivities of CME proteins in both cell types, we find interesting differences (Fig S14, S15). One major reason for several of the changes in sensitivity of the SB ratios is that 18 of the proteins in fibroblast have unknown copies. Algorithmically, these proteins can effectively absorb any ‘left-over’ interfaces without penalty. If they are assigned a high copy number, like CORO1A, which absorbs much of the binding to the extremely highly expressed ACTB (166 μM vs 2.4 μM in HeLa), then when it is removed, it acts as a strong disruptor to the network stoichiometry. This explains the sensitivity we observed for dynamins (DNM1 and DNM2), which are unusually unstable for such highly connected hub proteins—without CORO1A they are needed to bind to the millions of surplus actin copies. As a result, the correlation of sensitivity with node connectivity is much weaker in fibroblast than in HeLa cells (R=-0.07, Fig S16). Some of the strong pairwise couplings that emerge upon KD are also suppressed in fibroblast due to the unknown proteins. In HeLa, CAPZA2 was highly sensitive to CAPZB KD, but not in fibroblast (Fig 7C, red arrow). However, if CORO1A is removed from the network, we find that the strong coupling returns, demonstrating that these unconstrained nodes in the network help buffer against imbalances. Lastly, the co-factors of the chaperone HSC70/*HSPA8* are either not present (auxilin/DNAJC6=0) or unknown (GAK). Thus, while they were notably sub-stoichiometric in HeLa, their impact is not visible in the fibroblast cell.

### Synaptosome copy numbers and stoichiometry are relatively distinct from HeLa cell types

When comparing observed copy numbers between a neural synaptosome and HeLa, we find that they are only weakly correlated (R=0.05, Fig S17). Although the proteins we have classified as being highly expressed generally (light green, Fig 1) remain as the most highly expressed, most proteins have much higher concentrations in the synaptosome compared to HeLa (Fig S17). This is perhaps due to challenges in measuring very low copy numbers: the volume of the synaptosome is ^~^6000 times smaller than the HeLa cell^5^, so a single copy of a protein is already at 7 nM concentration, vs 1 pM in a HeLa. Relative to HeLa, the synaptosome CME network has also changed with the addition of AMPH, DNM3, SGIP1, SH3GL3, SNAP91/AP180, WAS, the cargo SYT1, and removal of TRIP10. Applying SBOPN, we observe that the stoichiometry in the synaptosome is thus distinctive from that of HeLa cells. When only considering proteins, the correlation is −0.07, with the addition of cargo it increases to 0.12 (Fig 7B), and with lipids, to 0.48. Unfortunately, in the synaptosome the copy numbers of many proteins involved in CME (49) have not been experimentally quantified, meaning that multiple nodes can act as buffers. This is seen to a lesser extent in the fibroblast cell. Nonetheless, we do find some sharp trends in SB ratios due to the overall increase in concentration of multiple proteins in the synaptosomes. When we evaluate SB ratios in the networks with just cytosolic proteins, or cytosolic protein +cargo (Fig. S18), we find that PIP5K1C, a lipid kinase that binds to AP-2 beta-subunit, is super-stoichiometric in synaptosomes (Fig 7B). In all other cells, kinases and phosphatases are sub-stoichiometric. Here, we see that the abundance of PIP5K1C has increased by a factor of over 900 in the synaptosome relative to HeLa, making it far more abundant than necessary for its partner AP2-beta appendage. This is true of not just PIP5K1C, and consequently, we see that the extent to which AP-2 is concentrated relative to its partners is greatly diminished in the synaptosome. Indeed, known protein copies in the synaptosome are within a factor of 10 of the AP-2 subunits, whereas in HeLa, proteins span a factor of ^~^1000 relative to AP-2. We also see some distinct trends in which proteins have sensitive or unstable SB ratios due to knockdown of others (Fig S19). DNM1 is now stable to perturbations in synaptosome (Fig 7D), unlike in HeLa cells, due to its dramatic increase in concentration levels, from 0.12 nM in HeLa to 16 mM in synaptosome, which exceeds the total DNM1,2,3 levels in HeLa (0.4uM) by a factor of 40.

Although we find that the SB ratios of the transmembrane cargo proteins themselves is similar in synaptosome and HeLa, the specific reason for VAMP2 and SNARE has swapped. VAMP2 has excessively high concentration: at 187 μM it exceeds any other cargo in any cell type, whereas it is present at ^~^0.008 μM in HeLa and fibroblast. SNARE is also highly concentrated at 0.7μM, but this is similar to the HeLa and Fibroblast. In the synaptosome, SNARE is now highly sensitive to knockdown of VAMP2, due to its imbalances (Fig. S19, S20). The VAMP2 and SNARE adaptor protein SNAP91/AP180, which is not present in HeLa or fibroblast, is also now the most concentrated adaptor protein, at 26 μM.

We can discuss changes in synaptosome stoichiometry in terms of the specific functional demands of the synaptosome cell type. VAMP2 is critical for the synaptic vesicle turnover, which is a primary role of the synaptosome. Distinct from the HeLa or fibroblast, the synaptosome is like a compartment at the end of a neuronal cell that contains mitochondria but not a nucleus^5^, and thus does not perform all of the same functions. Furthermore, at the synapse, there are multiple other forms of endocytosis or receptor uptake that are capable of faster (i.e. ultrafast endocytosis^83,89^) or internalization of larger sections of membrane (trogocytosis or phagocytosis), relative to CME. Unlike CME, these internalization pathways are not dependent on clathrin, and less so on AP-2. SYNJ2 and endophilin (SH3GL2), for example, both are important for ultrafast endocytosis^83^, not just CME; endophilin/SH3GL2 has increased concentration by a factor of 500, and SYNJ2 has increased concentration by a factor of 2400! It has nearly the same number of copies in the synaptosome as in the HeLa, despite vast differences in cell volume. Thus, the functional roles of these proteins has shifted substantially in the synapse, and even the neuronal isoforms of SYNJ2 and SYNJ1 have lost binding connections to the CME network (Table S2). SYNJ1, for example, no longer binds to AP-2.

### Across cell types, several trends are preserved regarding protein CME function

There are several trends in the abundance and stoichiometry of distinct protein classes across these 3 cell types. The proteins that assemble into rigid-like structures, actin (ACTB, ACTG1) and clathrin, are highly abundant, as are their respective disassembly proteins, cofilin (CFL1) and HSC70/*HSPA8*). Relative to the network of protein binding partners, these proteins are nearly always super-stoichiometric, or in excess supply (Fig 6, 7A–7B). In contrast, enzymes have low abundance, and are nearly always sub-stoichiometric, or in demand. We speculate that this is representative of more general control mechanisms throughout the network, where proteins with low abundance act as control nodes or gatekeepers that determine when or where assembly occurs. There is essentially a reservoir of clathrin and actin available to assemble, and a reservoir as well of their corresponding disassembly factors. Although these abundant proteins self-assemble, the clathrin interactions in particular are weak (>100 μM)^90^, and the clathrin-adaptor interactions are also typically weak^85^. High abundance supports a sufficient number of complexes even being formed, with the membrane 2D environment further stabilizing nucleation and growth^24^. With a reservoir of abundant proteins, the timing of these events is therefore controlled by essential co-factor proteins, as is the case for clathrin disassembly, which requires the auxilin or GAK proteins that have low abundance and sub-stoichiometry. Experimentally, it was shown that this clathrin disassembly machinery is found to localize to complete vesicles, not the pre-fission coated pits^91^. While sub-stoichiometry cannot ensure that co-factors will never localize to sites of clathrin-coated pits, by estimating the footprint of a single clathrin within a lattice as ^~^0.0009 μm^2^, even if 40% of the plasma membrane of a HeLa cell was coated in clathrin, more than half of all clathrin trimers would still be in solution. While HSC70 is sufficiently abundant to readily localize to coated pits, the sub-stoichiometric co-factors would help minimize disassembly at the plasma membrane.

For CME, we see a hierarchy of stoichiometries extending from the membrane. The highly connected lipids PI(4,5)P_2_ and PI/PS are also highly abundant and available for their many partners (although we note they both also have partners outside of the CME network). We then have the cargo adaptor proteins, which are mostly super-stoichiometric relative to their respective target transmembrane cargo. The packaging or selection of the cargo is then not limited by the number of adaptors, which are sufficient, but the competition of adaptor proteins with one another to assemble with AP-2 and clathrin. We see across cell types that the competition for binding to the hub interfaces on AP-2 and on clathrin drive many of their partners to be super-stoichiometric, or available in far excess of their accessible binding interfaces on AP-2 and clathrin. We also find that subunits of multi-protein complexes (AP-2, ARP2/3) tend to have similar abundances, consistent with experiment^21^, and exhibit more tightly linked stoichiometries. For example, the actin capping proteins and subunits of the ARP2/3 complex are in many cases sensitive to KD of their partners in all cell types. However, this is only true for subunits that connect to the larger network—protected subunits tend to be insensitive to perturbations across the network. Hence, while there are several distinctions that occur across cell-types as discussed above, because the network topology is very similar across cell types, we see several patterns of local and global disruption to stoichiometry that re-emerge in all cell types despite variations in protein abundances.

### Across cell types, several trends are preserved regarding abundance and topology

There are several trends in stoichiometry that arise due to the connectivity of proteins or their abundance levels. We expect these trends to persist across different network or processes, not just CME, as they reflect general optimality constraints for maintaining stoichiometry of binding throughout a highly connected network. Specifically, we find persistent positive correlations between SB ratios and abundance, with low copy number proteins, such as enzymes, being sub-stoichiometric, and highly abundant proteins being super-stoichiometric (Fig. 5B, S21). We also find that the most highly connected nodes tend to be stoichiometric, because it is often less costly in optimizing balance across the network (Fig. 5D, Fig S22). Several trends then emerge when we systematically knockdown proteins from the network and quantify the change in the SB distribution. Proteins that are most disruptive to the network stoichiometry upon KD are more abundant, and have higher connectivity, like AP-2 subunits (Fig. S8, S23). This is preserved across cell type, although the correlation is weakened by proteins such as dynamin1, which are highly connected but have low abundance in HeLa cells. In contrast, proteins that are sensitive and have unstable stoichiometry in response to KD tend to have few network connections, and lower copy numbers (Fig. 7C–7D, Fig S12, S16, S20). For example, DBNL exhibits an SB ratio that is unstable against a variety of perturbations, across all 3 cell types. DBNL has similar copy numbers across all cell types (0.25-0.8 mM) and only three connections, but they are all to highly connected hub interfaces. DBNL is an actin binding protein (ACTB and ACTG1) that also uses its SH3 domain to bind to the hub DNM1. For each cell type, we thus see that not only does removal of its immediate partners alter its SB ratio, but KD to many of the actin or DNM1 partners also alter its stoichiometry.

Most strikingly, proteins that are disruptive, like HSC70/*HSPA8*, are robust against perturbations to the network, appearing on the left hand sides of Fig 7C–D. This correlation between disruptiveness and stability against perturbations is stronger than either of the correlations of KD with copy number or connectivity, as it more directly compares how the stoichiometry of a protein impacts or is impacted by perturbations to the network (Fig S11, Fig. S24). An important caveat is that because the SB ratios and sensitivities depend on the protein network, for proteins whose KD has a minimal impact in CME, it could be more impactful if that protein functions more prominently in other biological pathways. We included key cytoskeletal proteins for CME in our network^12^, but not an exhaustive network of proteins that impact actin assembly. Relative abundance can also be shifted with additional network components, as we see most clearly with the lipids (Fig 6B), but HSC70/HSPA8 is another example of a protein with additional cellular functions and binding partners outside of this network that could reduce its availability to the CME proteins if they share the same binding interface^8^. Lastly, we note that the presence of unknown protein copy numbers in the network can act as sinks to absorb binding interactions with abundant proteins. These network nodes then tend to suppress couplings, as we saw in the fibroblast when CORO1A was removed from the network. The more experimental copy numbers that are available, the more reliable and predictive we expect the trends and patterns to be.

## Conclusions

The analysis we have performed here on our newly constructed interface-resolved CME network integrates structural data, biochemical studies, and protein abundances to reveal patterns of stoichiometry that directly connect to protein function. One of our more surprising findings is how proteins that are quite similar functionally, like the cargo adaptor proteins DAB2 and LDLRAP1/ARH, can nonetheless display varying disruptiveness upon removal. These results would be difficult to intuit, but in the context of the stoichiometric balance model, they can be explained by differences in copy number and connections to distinct components of the network, with DAB2 being the most unusually connected adaptor protein.

The stoichiometric relationships between proteins we observe are not possible with just the network or the copy numbers alone. Clathrin is highly abundant and super-stoichiometric, but this is not because it is in fact more numerous than all its binding partners—it is not. However, when its binding partners are adjusted for whether they are sub- or super-stoichiometric themselves, or occupied with other binding partners, the final balanced copies of clathrin are still in excess supply. This theme re-appears for other proteins as well: just because a protein (like clathrin or PICALM) appears to be sub-stoichiometric relative to a set of partners, does not mean that it is sub-stoichiometric overall. Instead, the partners can be super-stoichiometric due to their other connections. Further, although some protein couplings emerge between direct partners, several of the sensitivities to computational knockdown result not from immediate partners, but via competition for proteins that share binding to a hub interface (e.g. DAB2, DBNL). Although the stoichiometric analysis performed here does not account for the strength of the binding interactions between proteins, effectively assuming they all have infinite strength, the SBOPN method can be generalized to introduce weights to interactions to mimic weak vs strong binding. Hence the relative stoichiometry of the interactions can be re-assessed in light of the need to have highly abundant proteins, for example, to ensure any complexes are formed for weak interactions. Because many domain-domain interactions also share similar affinities to one another (such as several AP-2 appendage and SLIM motif interactions being ^~^μM strength) the assumption of identical edge weights for each hub interface is, at least, reasonable to motivate predictions and describe trends. This analysis is not able to report on the spatial or temporal regulation of CME, as it is based on a steady-state population of bound structures. CME is, of course, sensitive to spatial localization to the membrane, and although stochastic, has an average timescale of ^~^ 1 minute to proceed, out-of-equilibrium, from cytosolic components to a budded, clathrin-coated vesicle. It also is dependent on mechanical forces required to bend the membrane. However, the components in our model are constitutively expressed, so we do expect these proteins to be present in the cell simultaneously. Spatiotemporal dynamics with mechanics cannot be tractably studied in the context of a network of this size, with almost 100 distinct components, and performing KDs of one or many proteins systematically introduces further expense. While modeling in cell biology is reaching closer to achieving these types of simulations^62^, our approach here is a powerful complement that requires no parameterization, while still yielding meaningful insight. Several of the trends we observe also make intuitive sense, such as highly abundant and highly connected proteins being more disruptive to the SB distribution. Abundant proteins and hub proteins are often more essential^92 93^.

Critically, our SBOPN method is based on a physical model of binding specificity and competition, and thus we can trace the origins of all SB ratios we observe, making outputs predictive and not purely correlative. Because the SB ratios report on the ability of each protein to form complexes, changes in the SB ratio define how a new set of interactions and complexes will form upon knockdown, which would alter the dynamics. The exact extent to which complexes will change upon knockdown depend on binding strengths and dynamics, but our results predict that an experimental knockdown of DAB2, for example, would be significantly more impactful in affecting CME dynamics in HeLa cells than in fibroblast, as it is much more abundant and out-competes other adaptors in binding to AP-2. A recent experimental study demonstrated how single KD of multiple endocytic adaptor proteins in epithelial cells impacted clathrin coated structure formation, finding a surprising lack of correlation between the impact on early CME stages and the amount of bulk transferrin uptake^88^. Our analysis further suggests where specific changes might be expected to occur on a protein-by-protein basis, made possible by our analysis of every protein, lipid, and cargo response to perturbations to the network. Our analysis can also be used to perform ‘over-expression’ of nodes, or more minor perturbations than occur upon complete removal of a protein from the network.

The curated network we have constructed provides a foundation for further evolutionary or dynamical systems studies. With binding interactions resolved down to the interface or domain, the changes in specificity of individual binding interactions can be compared throughout evolution^59^, across diseases, and changes in network connectivity can be compared against the interface-resolved network in yeast^59,65^, for example. The CME interactome here may be incomplete, as unresolved interactions or as-yet-unknown proteins may play a role. Nevertheless, we know from our analysis that the addition or removal of nodes (or interactions) from the network can have minimal effects on stoichiometry, or it can indicate important correlations. Thus, we expect the conclusions from our analysis to provide a robust foundation from which to build in additional evidence of CME interactions or proteins.

## METHODS

### Network Construction

To construct the mammalian CME interface-resolved network, we initially considered 86 proteins. These 86 cytoplasmic proteins were selected due to their functional importance to the endocytosis pathway based on a systems-level study^10^, a comprehensive review ^11^, as well as homology to proteins in the yeast interactome^65^. To begin constructing our interface-resolved PPIN, we pooled unique PPIs mediated by only proteins listed in our original 86-protein CME list, downloaded from 3 open-source online molecular interaction repositories compiled from comprehensive curation efforts: Biological General Repository for Interaction Datasets^63^, IntAct^94^, and Mentha^95^. BioGrid provided the highest number of reported PPIs. IntAct is the only database that also curates information on domain resolution when possible.

Our completed interface-resolved human CME PPIN contains 82 proteins and 617 edges. Four of the original proteins either lacked reported PPIs in databases or lacked domain resolution for interactions. Of the 617 edges assigned, 433 were identified by a repository and interfaces were assigned with sufficient literature evidence, and 184 edges were added by manual curation, primarily due to studies on interactions with highly homologous proteins from other mammals (see Table S1 for % homology). We removed 31 edges due to suspected indirect and/or false interactions, and 202 edges downloaded from the repositories were left unassigned due to insufficient literature evidence to define binding sites. Overall, our filtered interface-resolved mammalian CME PPIN contains 82 proteins, 5 phospholipids, 9 transmembrane cargo (receptor/SNARE) groups that include 31 unique cargo proteins, and 396 distinct interfaces across all proteins and lipids. Our transmembrane cargo groups are YXXØ (where Y is tyrosine, Ø is a bulky hydrophobic residue and X is any residue), GPCRs, low density lipoprotein (LDL) receptors/phosphotyrosine binding domain targets (PTB), ubiquitylated cargo, specific VAMPs and a broader class of SNAREs, as their adaptor partners are known or inferred from sequence homology, with the complete list and classification in Table S1.

Below, we define the protocols and rules we used for delineating interface-resolved PPIs, given sufficient data, and rules for distinguishing interfaces within overlapping binding regions (Figure S1). Domains that mediate a PPI could be assigned based on varying levels of evidence, listed in descending order of definitiveness: both identified via co-crystal structure; both identified with residue information derived from in vivo and/or in vitro experiments; one identified and the other has been inferred from mammalian or yeast homology; one identified with supporting in vivo and/or in vitro evidence, and the other speculated; both inferred; and both speculated. The most straightforward assignments are based on crystal structures of the human proteins in co-complex with one another, with the majority of assignments, however, being based on biochemical methods between either the human proteins, or mammalian and yeast homologs.

#### Assigning interfaces to PPIs

We document relevant details, e.g. PubMed IDs corresponding to studies sourced, and justifications for binding interactions in Table S3, as we individually annotate each PPI pulled from the repositories. We kept two separate lists of mammalian (including *Bos taurus, Gallus gallus, Mus musculus,* and *Rattus*) and yeast homologs for each protein, along with their percentage values of sequence identity relative to their human counterparts determined from BLAST^96^. We collected homologs that reached a sequence identity score ≥60%.

We constructed a decision tree which we iterated through for every PPI per protein in constructing our interactome (Fig S1). In brief, we first chose a protein and pooled its unique PPIs across the 3 databases. Starting with one of its PPIs, our assignment task starts with a question that asks for evidence of highest specificity derived from a co-complex crystal structure, if available. Otherwise, the decision-making process continues with a series of questions that descends in orders of definitiveness, from residue level experimental support to: 1) whether domain information is available for both proteins or either protein; 2) whether domains for both or one of the proteins could be inferred from yeast or mammalian homology referring to our lists of BLAST scores; 3) whether domains for both or one of the proteins could be speculated; and 4) if neither domain could be speculated, whether literature support for PPIs were derived from high-throughput (HT) methodologies such as affinity capture-mass spectrometry (AC-MS)/MS and yeast-two hybrid (Y2H) assays. Hence, this process required substantial manual curation of the literature.

In addition to careful annotation of each PPI, we kept track of all the domains per protein. In order to define interfaces for an interaction, we either had to select one of the interfaces we had determined so far or add a new interface. We began with interface assignment with those determined from SMART^64^. In some cases, these domains mediated more than one PPI, hence containing multiple binding sites based on residue subsets (see Section: Splitting PPI binding domains into multiple subdomains, below). Interfaces could thus be broadly classified as structured (or globular “Glob.”) domains, specific residues within a structured domain, or short stretches of residues that were primarily part of unstructured regions, which we labeled Short LInear Motifs (SLIMs)^71^. In the process, we actively updated the domains of the partner proteins.

#### Matched edges

Of the 617 edges in our PPIN, 443 are marked “MATCHED”. These interactions were identified by at least one of the 3 repositories and were retained in the network given literature evidence supporting a direct and specific interaction.

#### Splitting binding domains into multiple binding interfaces

Specific binding sites and interfaces often lie in large, spanning regions that define an entire folded domain. These domains were split if residues have been reported to mediate the interaction. We do this because partners for a spanning region of a protein can be noncompetitive, meaning both interactions can simultaneously be formed, and because not all residues are important for specificity of the interaction. We do not verify that all distinct interfaces are sterically accessible simultaneously, as done in one study^27^, focusing instead on the specificity of the individual interface residues for distinct partners, similar to previous work^65^. PRDs provide an example of unstructured regions containing multiple sequential binding motifs with often distinctive specificity for binding partners^97^. For example, the Huntingtin-interacting protein 1 (HIP1R) has a PRD including residues 1016 to 1068. The HIP1R interaction with cortactin (CTTN) specifically requires residue 1016, whereas mutations of essential residues in the 1025-1030 motif do not seem to impact binding. Residues 1025-1030 mediate specific binding of HIP1R to the three SH3 domains of SH3KBP1/CIN85, hence these regions are designated as distinct interfaces. BAR domain proteins provide an example of a structured domain with multiple binding interfaces. These domains can simultaneously form dimers with one another, bind to the membrane, and form higher-order oligomers. Thus, for each BAR domain, we created separate interfaces for dimerization, lipid binding, and oligomerization.

#### Specificity for distinct copies of repeated domains

Similar to the PRD example above, proteins also can contain repeated copies of a structured domain that have distinct specificity, requiring us to list each copy as its own interface. As an example, intersectin 1/2 (ITSN1/2) proteins each contains 5 unique SH3 domains, SH3A-E, that have distinct binding specificities to protein partners such as DNM2 and WAS.

#### Speculating interfaces for binding interactions

Many proteins in our CME list share the same protein binding partners and sometimes have shared binding regions. We use this information, coupled with homology, to define interface-resolved interactions into our PPIN, despite not having direct evidence of the specific domains used for the binding pair. To demonstrate our definition for speculating interactions, we use A1, A2, A3 isoforms of endophilins (SH3GL2/1/3 respectively), along with AMPH and BIN1, as they all share similar domain architecture and overlap in binding partners, to help define binding interactions to DNM1 and DNM2. We use homology to define SH3-PRD interactions that are known to DNM2-SH3GL1 for speculating binding interfaces for DNM1-BIN1 and DNM2-BIN1.

#### Added edges

In our curated PPIN, 184 new edges were added to account for interactions that were not present in the human PPI databases, but were supported by other literature studies. Of the 184, 80 of these added edges are lipid and cargo interactions, which we did not initially pull from the repositories. Most of the other added edges were not identified by any of the 3 repositories and were added during our re-evaluation of “MATCHED” PPIs via literature research. We sought to use homology for resolving interfaces of added PPIs, which we make note of below.

#### Using homology to add functional PPIs based on definitive biochemical evidence

Some interface-resolved “ADDED” PPIs were included using homology-based inference. These are interface-resolved PPIs informed by low-throughput biochemical experiments using homologous proteins. We take DBNL-DNM1 as an example. Mouse Actin-binding protein 1 (mAbp1), physically associates with rat Dnm1, serving to be a physical link between the actin cytoskeleton and endocytosis in both membrane transport processes in neuronal and non-neuronal cells at actin-rich sites, confirmed by immunoprecipitation and immunofluorescence microscopy within the same study. Additionally, because mouse Abp1 and human Dbnl share a sequence identity score of 85.39% and rat and human Dnm1 a high score of 98.03%, we thus added DBNL-DNM1 into our network.

#### Using homology to infer interfaces for dimerization and oligomerization

Since CME depends on dimerization and oligomerization of BAR-containing proteins such as F-BAR domain only proteins 1/2 (FCHO1/2), these PPIs were added into our network, if not already identified. For example, in our network, we identified FCHO2:FCHO2 as a “MATCHED” edge, but not FCHO1:FCHO1 nor FCHO1:FCHO2. However, the crystal structures of the yeast homolog SYP1 for human FCHO1/2 proteins and the structure of FCHO1 paralog FCHO2 homodimer have both been resolved. Using this information, we inferred interfaces for homodimerization and oligomerization interactions for FCHO1, adding the FCHO1:FCHO1 and FCHO1:FCHO2 PPIs into our network.

#### Inclusion of protein-lipid and protein-cargo interactions

Unlike other existing protein-protein interactomes, our interface-resolved network integrates relevant CME protein-lipid interactions as well as protein-cargo interactions (Table S3). Our inclusivity of lipid-binding interactions builds off of known identification of lipid-binding domains such as BAR, ENTH, and PH domains, including 42 of our proteins. Some of these interactions are enzymatic reactions that help regulate the population of phosphoinositides at the plasma membrane. For cargo interactions, although our interactome is not expected to be comprehensive in including all receptor cargos, we drew from reviews that classified types of cargo interactions specific to CME^70^.

#### Unassigned edges

202 edges were left unassigned from our network. High throughput Y2H- or AC-MS/MS-based screening used to construct large integrated protein interaction libraries PPIs can produce false positives and false negatives. If we could not find additional evidence to support an interaction between two proteins, this PPI was unassigned. Interactions that were unassigned were also those pulled from low-throughput experiments that lacked domain resolution.

#### Removed edges

31 edges were marked removed from our original list, as there was evidence from additional experiments that they were indirect or false. As reported in the construction and characterization study for the yeast CME interactome^65^ the PDB structure of the ARP2/3 complex^98^ was used to assign interfaces, and thus all other subunit contacts found in the 3 repositories were removed. 11 of the 31, or 35.5% of the removed edges, were Arp2/3 subunit interactions disregarded for this reason. PPIs were also removed if they were reported in a high-throughput study but proteins were biochemically shown to not interact. Phosphatidylinositol-4-phosphate 5-kinase type 1 gamma interacts with the β appendage of AP-2, not the α subunit (Table S3).

#### Copy numbers by cell type

Organisms contain many different cell types, each of which produces its own pattern of protein expression levels and in some cases distinct splicing isoforms of protein encoding genes. Our study collected copy numbers from rat synaptosome ^5^, mouse fibroblast^4^, and two human HeLa cell ^2,3^ studies. In all three cell types, some of our proteins had no copy numbers reported. Proteins whose copy numbers were not reported in a study might be due to low natural abundance with expression levels undetectable. Therefore, for these proteins we used the Human Protein Atlas^6^ and Proteomics DB^7^, which report protein expression levels by tissue types, to help determine whether proteins with no reported copy numbers had unknown copy numbers, or zero copy numbers. Specifically, we looked for expression for the synaptosome proteins based on neuronal tissue (brain, cortex); for the fibroblast proteins based on bone marrow, stroma, soft tissue and skin fibroblasts; and for the HeLa proteins based on cervix, uterine, and squamous epithelial cells. Proteins with zero expression were excluded from subsequent stoichiometric balance analyses by removing the proteins from the network. If the databases indicated any expression level of a protein in a specific cell type, it was left in the network. Proteins that do not have constraints on copy numbers (“—“) thus can constrain the stoichiometric balance of their partners, but they are not penalized at all for deviating from their observed (unknown) copy number.

### Stoichiometric balance optimization

The Stoichiometric Balance Optimization of Protein Networks (SBOPN) algorithm has been described previously^23^. Briefly, the algorithm requires the network of interacting proteins, with interfaces resolved on their parent proteins. The solution of the number of complexes for each pair of binding partners can be formulated as the optimal of a quadratic function, with linear constraints. Specifically, the copy numbers of each interface are defined as the sum over all complexes it is part of. Two soft constraints are applied. First, for a protein with multiple interfaces, there is a penalty to making them highly distinct from one another. Second, a protein’s copies can be constrained to a target value, typically the observed copies of a protein. There is one parameter, *α*, which controls which of these two soft constraints is more tightly applied, where a low value of *α* forces all interfaces to be the same within a protein. We use here a value of *α*=1, which allows fluctuations in values across interfaces, similar to known variances in measured copy numbers. Given the optimal number of complexes that are solved for, we can then calculate the balanced number of interface and thus protein copy numbers. The code is available on https://github.com/mjohn218/StoichiometricBalance, along with input files for this network and knockdown simulations. The interactive network file generated using Cytoscape v3.7.2^99^ is available as part of supplemental material.

Pearson correlation coefficients (R) between distributions were always applied to log10 values of concentrations, and to log10 values of SB ratios. The distances between SB distributions was calculated using the Jensen-Shannon distance, where the distributions *p* and *q* were first normalized:

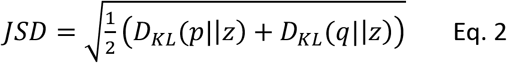

where 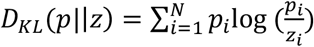 is the Kullback-Leibler divergence, and 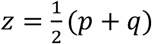. The sum loops over all proteins *N* with known copy numbers. If we used a chi-squared distance instead of the JSD, the results were similar. The sensitivity or stability of a protein’s SB ratio to removal of nodes from the network was calculated for each protein *p* as

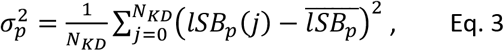

where 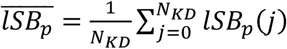, *N*_KD_ is the number of computational ‘knockdowns’ or node removals for a network, and *lSB*_*p*_(*j*) is the log10(SB) of protein *p* for a ‘knockdown’ *j*.

## ACKNOWLEDGEMENTS

We gratefully acknowledge funding by an NIH MIRA R35GM133644 to MEJ. MH was supported through an AMGEN scholarship. DH was supported by the intramural research program of the NIH.

