## Supplementary Figures and Tables for "Integrating protein copy numbers with interaction networks to quantify stoichiometry in mammalian endocytosis"

For

**Supplemental Figures S1-S24**

**Supplemental Tables S4-S6**

### Supplemental Figures

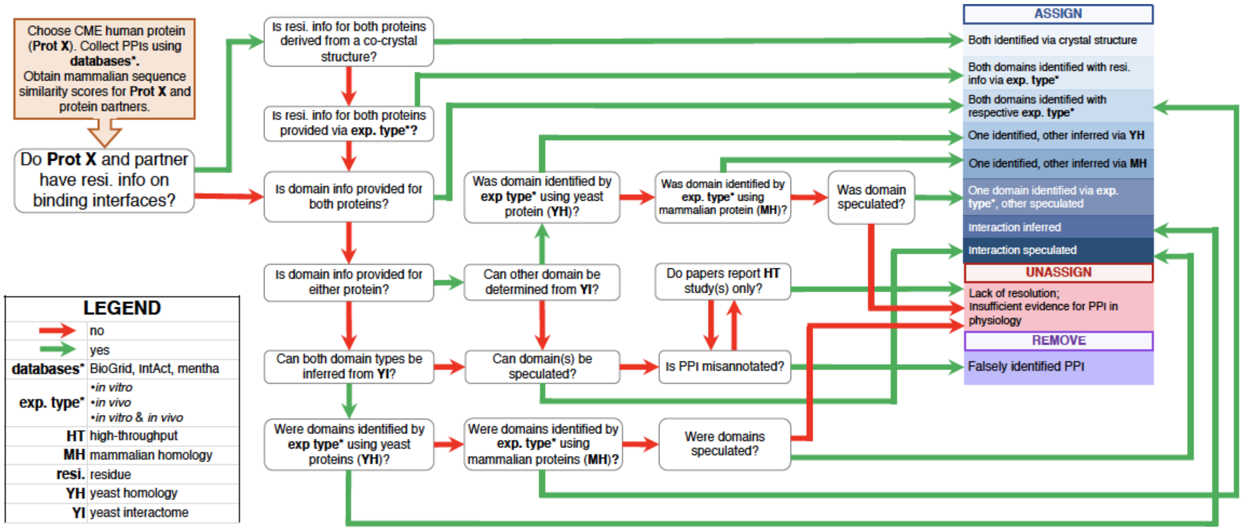


**Figure S1**. Decision-making workflow to annotate the assignment of a domain-resolved PPI based on experimental resolution. The decision-making series is followed by the network curator to determine whether the PPIs downloaded directly from BioGrid, IntAct, and mentha will be retained in the resulting CME network. There are interactions that differ in resolution based on the accuracy of the experimental methodology used to elucidate domain-resolved PPIs, in which case the decision-maker will delineate by listing specific experimental types used to elucidate them. With interactions that lack resi. and/or domain information, the user will be asked to infer domain partners of respective proteins using the YI curated by Johnson and Hummer (2013) with user's annotation detailing his/her inference of interacting domains based on YH. There are interface-resolved PPIs present in YI that have also been inferred using MH, in which case, depending on experimental type and whether domain information is provided for one or both proteins, the user will annotate with the appropriate level of experimental certainty, utilizing the provided BLAST sequence similarity scores of the human protein and its mouse/rat homologs. Interactions are speculated if no residue nor domain info is listed, but PPI is suspected to be mediated by two interfaces based on the domain architecture of the proteins and whether a similar interaction type is present in the CME network based on function. An interaction is unassigned if domain information is lacking. An interaction is removed if the PPI has been incorrectly annotated in the databases and is not physiologically functional.
**exp. type**: experimental type; **MH**: mammalian homology; **resi.**: residue; **YH**: yeast homology; **YI**: yeast interactome.

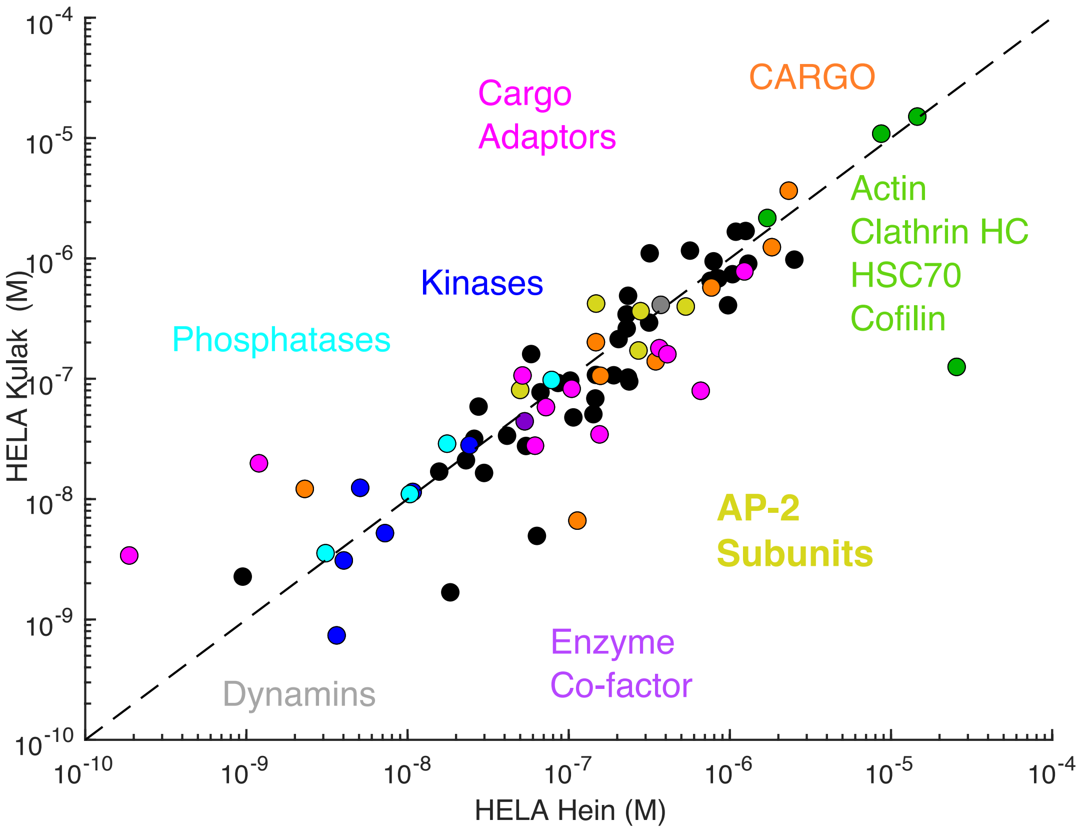


**Fig S2: Correlation is high between CME protein copy numbers in two HeLa cell studies.** We note that correlation is applied to the log10 values of the copy numbers, to identify correlations by order-of-magnitudes, giving R=0.83. Otherwise, on a linear scale, correlations are dominated by deviations between highly expressed proteins (C=54). The notable outlier is ACTG1, which differs by a factor of ~200 between studies. The dashed line indicates 1:1 correspondence (slope=1). In the main text, we use the average values. Proteins that were observed in only one study are not shown.

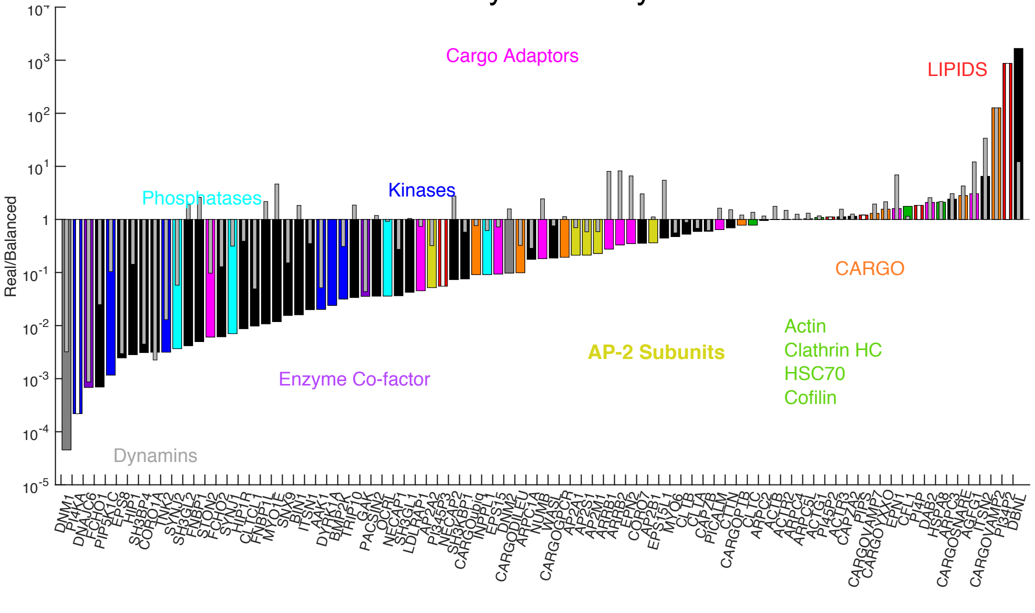

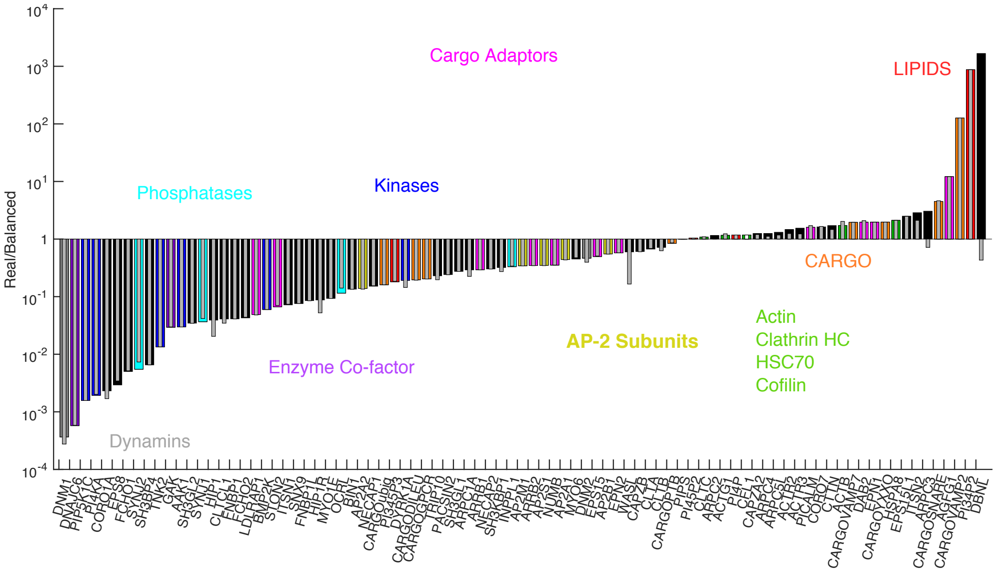


**Figure S3. SB ratio in HeLa cells, network with protein, cargo and lipids.** Top) The colorful bars indicated stoichiometry with lipids present, and the gray bars indicate the shift in stoichiometry after all the lipids are removed. After lipid KD, the super stoichiometric proteins are still super-stoichiometric, and the stoichiometric proteins have shifted to super-stoichiometric. Sub-stoichiometric proteins are less sub-stoichiometric, and several of them, most notably the cargo adaptors (pink) have switched to being super-stoichiometric. Bottom) The SB ratios are similar when we change the protein: lipid stoichiometry, but the cytosolic proteins are shifted to being less sub-stoichiometric. For proteins binding to PI/PS, we change to stoichiometry from 1:1 to 1:10, given the larger footprint of lipids they can bind to. For PIP2, we change the stoichiometry of AP-2 from 1:1 to 1:2. Gray bars indicate changes due to a CFL1 knock-down.

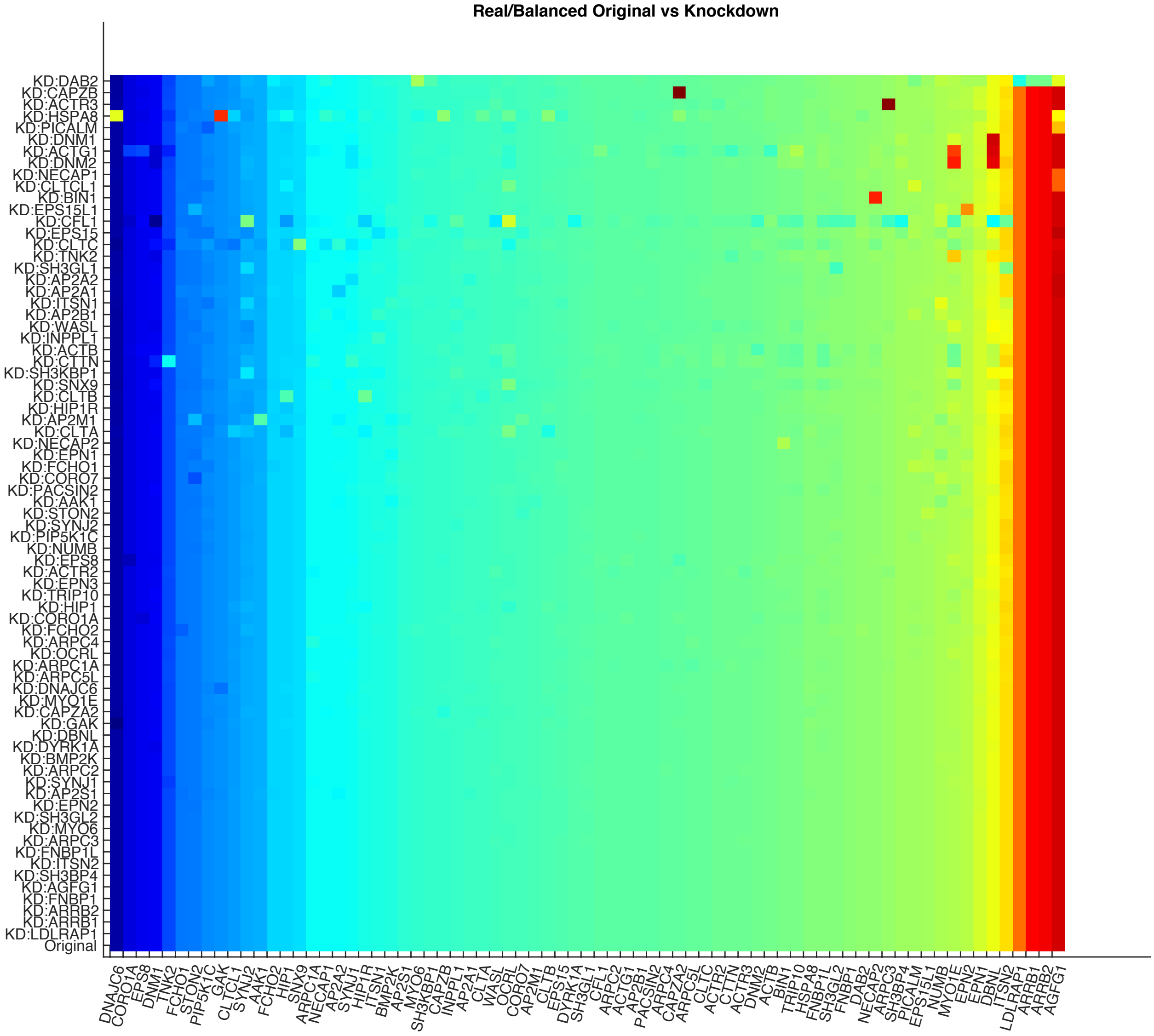


**Fig S4. SB ratios change when nodes are removed or 'knocked-down' (KD) from the network, with most impactful KDs at the top.** These SB ratios are sorted along the x-axis from smallest to largest in the network of cytosolic proteins (bottom row). We see in the top row that KD of DAB2 has a broad global effect on the SB distribution. In contrast, KD of CAPZB has a highly localized effect on CAPZA2. KD of the adaptors LDLRAP1, ARRB1 and ARRB2, at the bottom, have almost imperceptible impact on the network SB distribution.

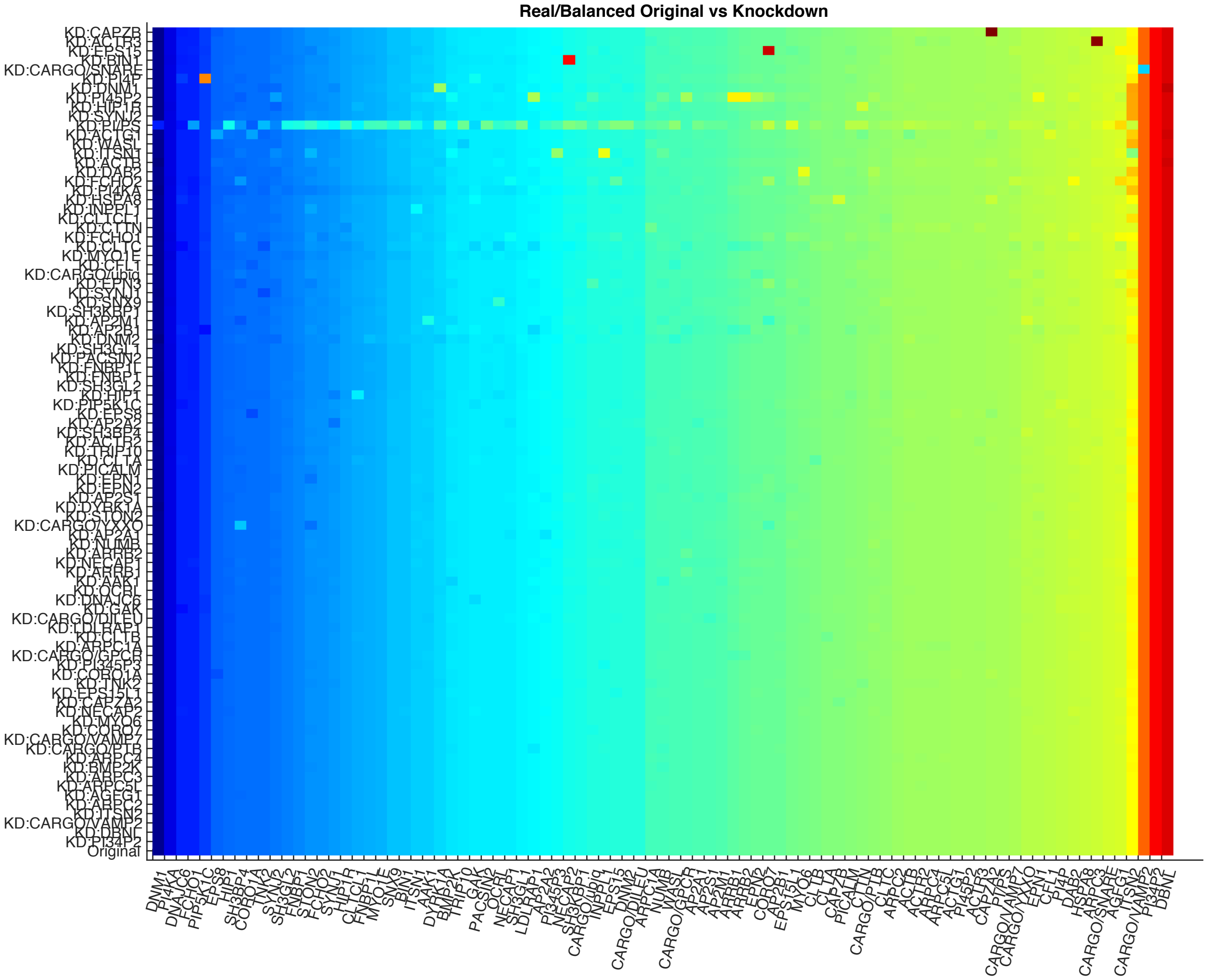


**Figure S5. SB ratios change when nodes are removed or 'knocked-down' (KD) from the network with proteins, cargo, and lipids.** These SB ratios are sorted along the x-axis from smallest to largest in the full network of cytosolic proteins+membrane+cargo (bottom row). We see in the top row that KD of CAPZB has a highly localized effect on CAPZA2. The KD of either PI45P2 or PIPS cause broad global changes across the SB ratios. PI4P is more localized, and the KD of minimally connected PI34P2 and PI345P3 have minimal impact.

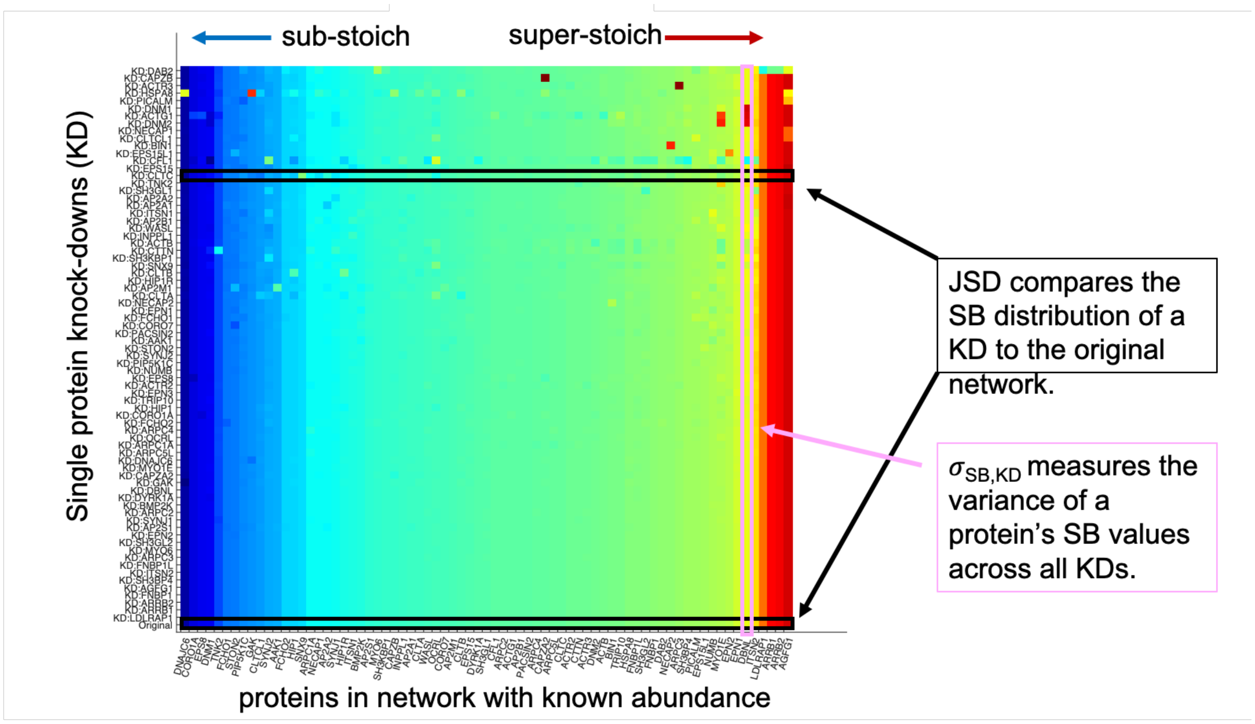


**Figure S6. Analysis of the matrix of stoichiometry balance (SB) ratios.** This matrix can be used to quantify the Jensen-Shannon distance (JSD) between the SB distribution of a KD relative to the original network (Eq 1 main text). It can also be used to quantify the sensitivity of a protein to KD, or $\sigma_{SB,KD}$ which reports on how unstable a protein's SB ratio is (Eq 2 main text).

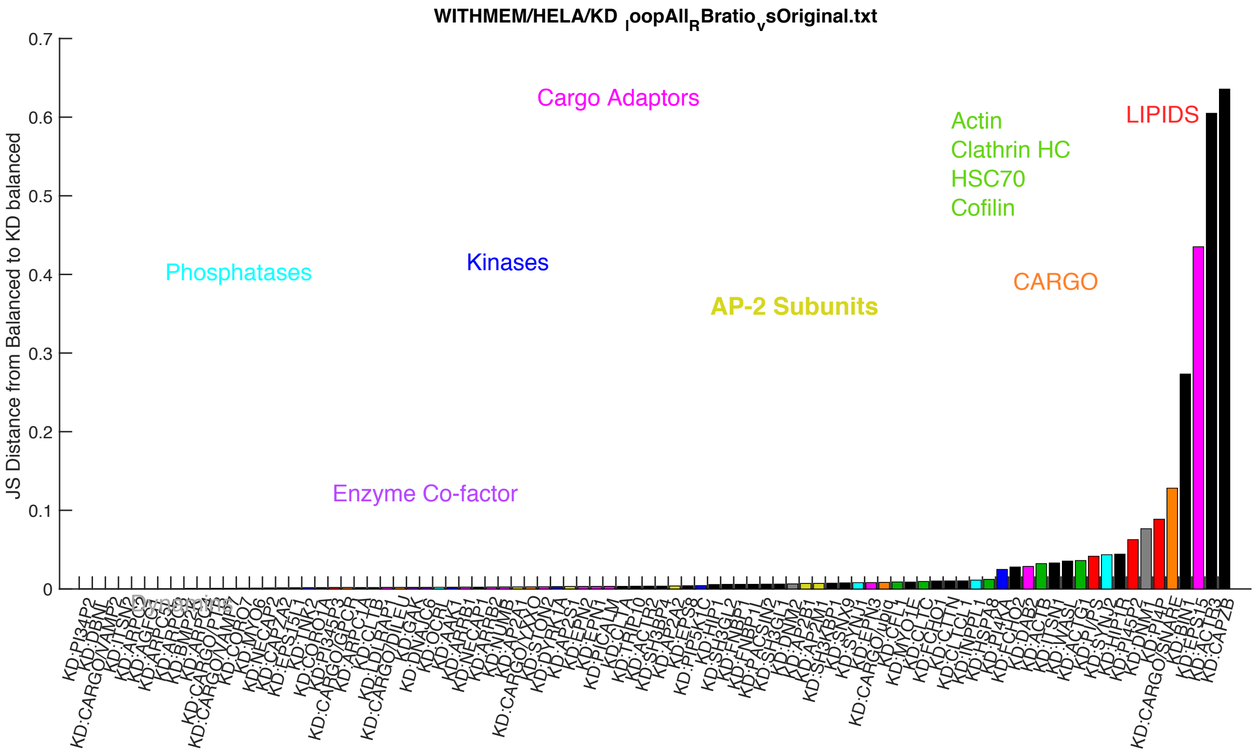

**Fig S7. Single protein KDs sorted by the Jensen-Shannon Distance (JSD) between SB distributions before and after KD.** Proteins on the right are disruptive to the SB distribution of the network. Proteins on the left have minimal impact on the SB distribution of the network. Results from the network with protein+cargo+lipids.

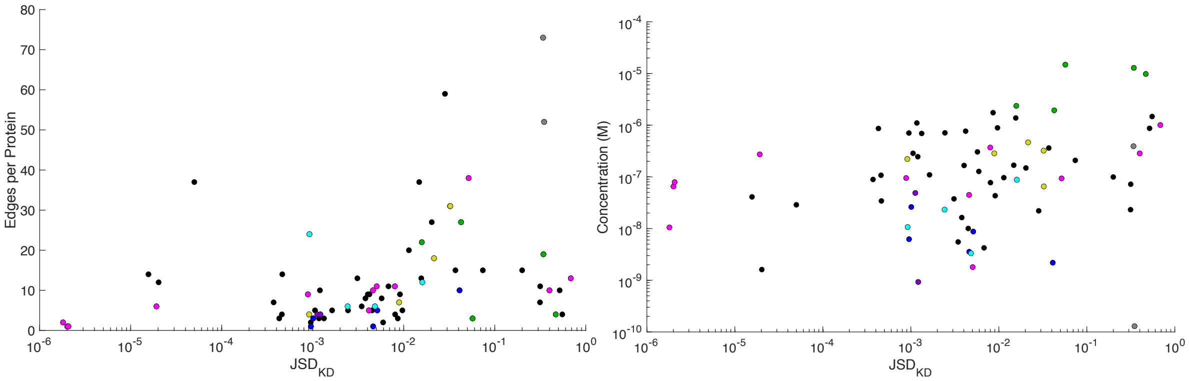


**Fig S8. The impact that a protein's KD has on the network stoichiometry is correlated with the connectivity of the protein, and its concentration.** The JSD for each protein's KD is plotted against number of edges on the protein (left), with R=0.37, and with the protein abundance, R=0.27.

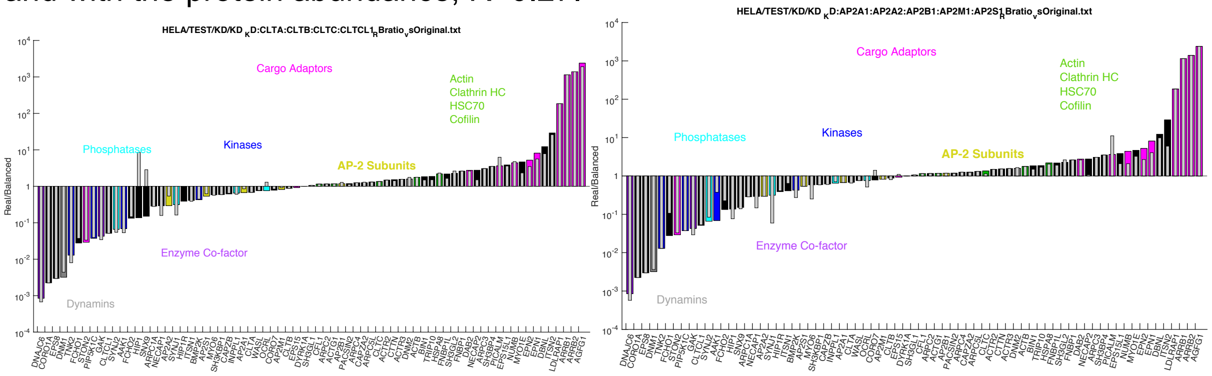


**Fig S9. Knockdowns of all clathrin trimer chains and all AP-2 subunits.** Left side is the clathrin chains, which affects the AP-2 subunits, some adaptors, and HIP1 and SNX9. Right hand side is AP-2 subunits knock-down, which affects several adaptors, AAK1 and SYNJ1.

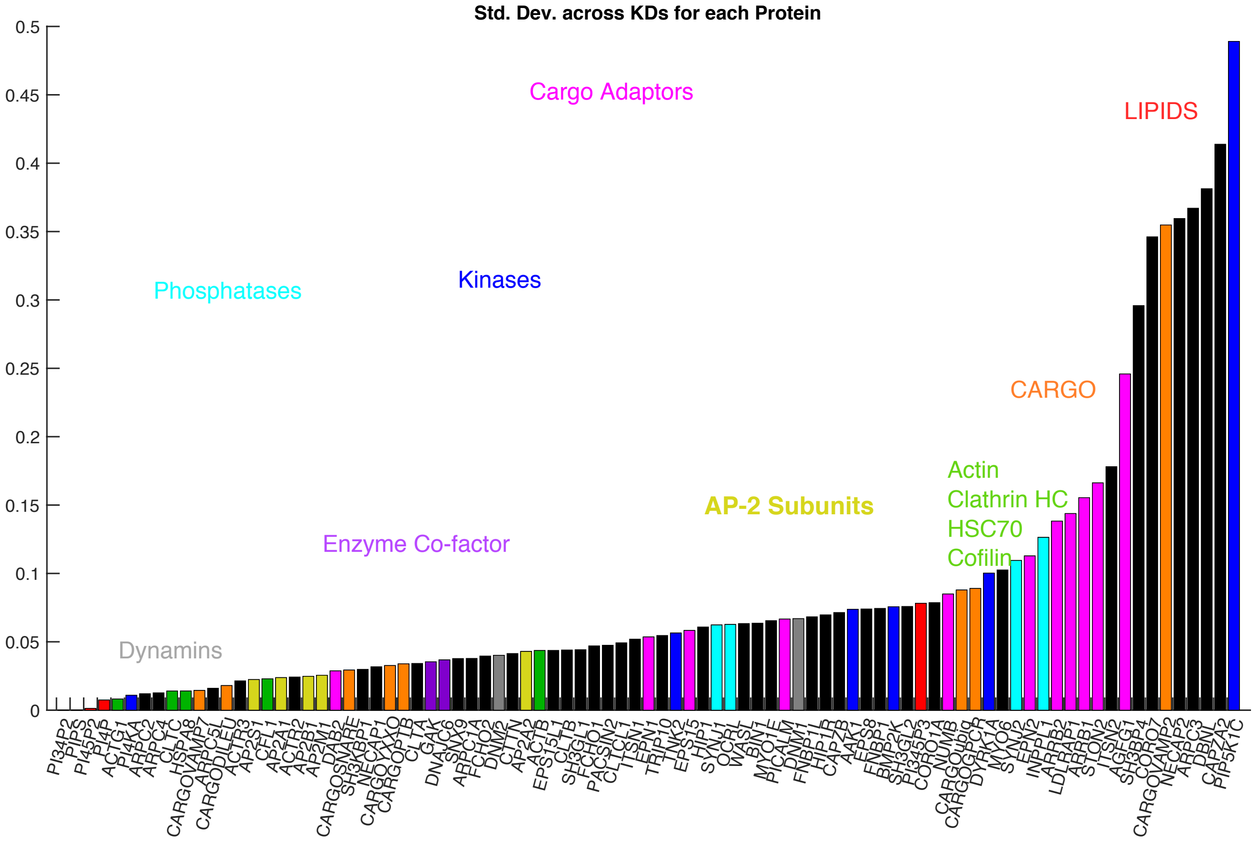


**Figure S10**: **Sensitivity of each protein's SB values to knock-downs, $\sigma_{SB,KD}$.** Sensitive or unstable proteins are on the RHS, their SB value changes as various other proteins are removed from the network. Robust proteins are on the LHS, they have SB ratios that are stable when other proteins are removed. Standard deviation is performed on the log10 SB values. LDLRAP1 is the only cargo adaptor where we see that the adaptor stoichiometry is determined by its cargo, rather than the cargo stoichiometry being determined by the more highly connected adaptors. LDLRAP1 switches to sub-stoichiometric: it has quite low copies (0.01 mM) and binds to a highly expressed cargo, the phosphotyrosine-motif NPXY found in LDL receptors (called the cargo_PTB) (0.67mM), which needs more copies of LDLRAP1 than are available.

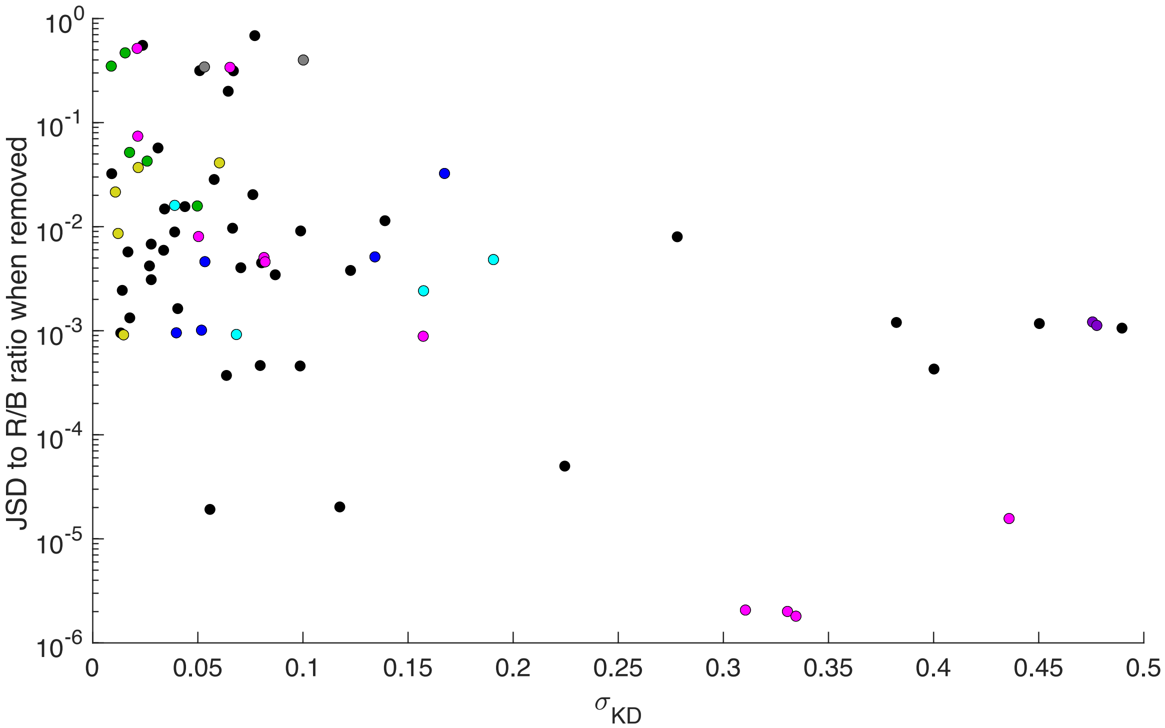


**Figure S11. The sensitivity of each protein is anticorrelated with its disruption to others.** Hela cells, Disruptive proteins tend to have stable SB ratios. Correlation is R=-0.51, applied to log10(JSD) vs $\boldsymbol{\sigma_{SB,KD}}$. With the membrane, R= -0.1827.

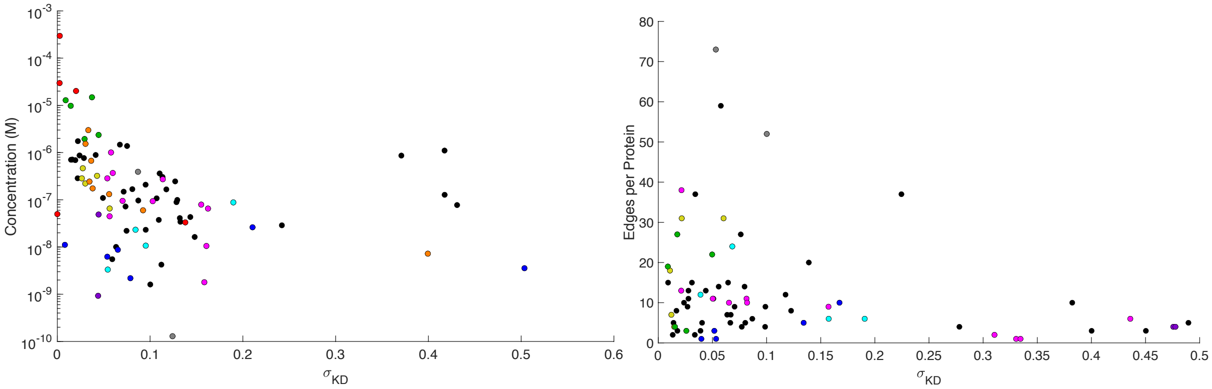


**Figure S12. Sensitivity of SB ratios to knock-down correlates with concentration and network connectivity.** Left, we see two regions: proteins with lower sensitivity have strong correlation with concentration. Unstable proteins do not correlate with concentration though, and overall R=-0.29 with lipids, and -0.19 with just proteins. Right) Compared to edge connectivity, we see that unstable proteins have low connectivity. Stable proteins vary, but highly connected proteins tend towards higher stability. R=-0.23. With lipids, R=-0.1.

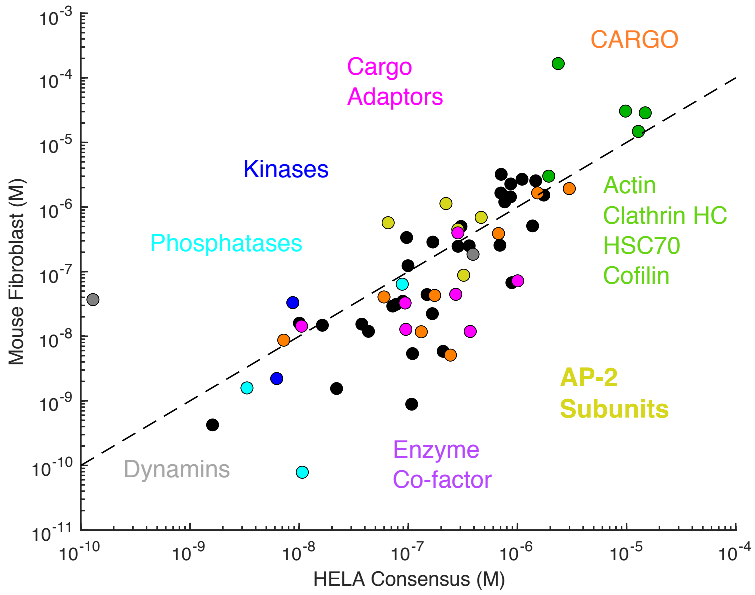


**Fig S13. Correlation between protein concentrations of the Mouse Fibroblast and the Hela cells.** On the log10 scale, R=0.77. An outlier is ACTB, which is 120M copies (166uM) in Fibroblast vs 2M (2.4uM) in HELA. Proteins with known abundance in only one cell type do not appear. Unknown proteins in fibroblast are ARRB1, AMPH, CORO1A, CORO7, DYRK1A,FCHO1, FNBP1, GAK, ITSN2, MYO1E, NUMB, OCRL, PI4KA, PIP5K1C, SGIP1, STON2, TNK2, WAS.

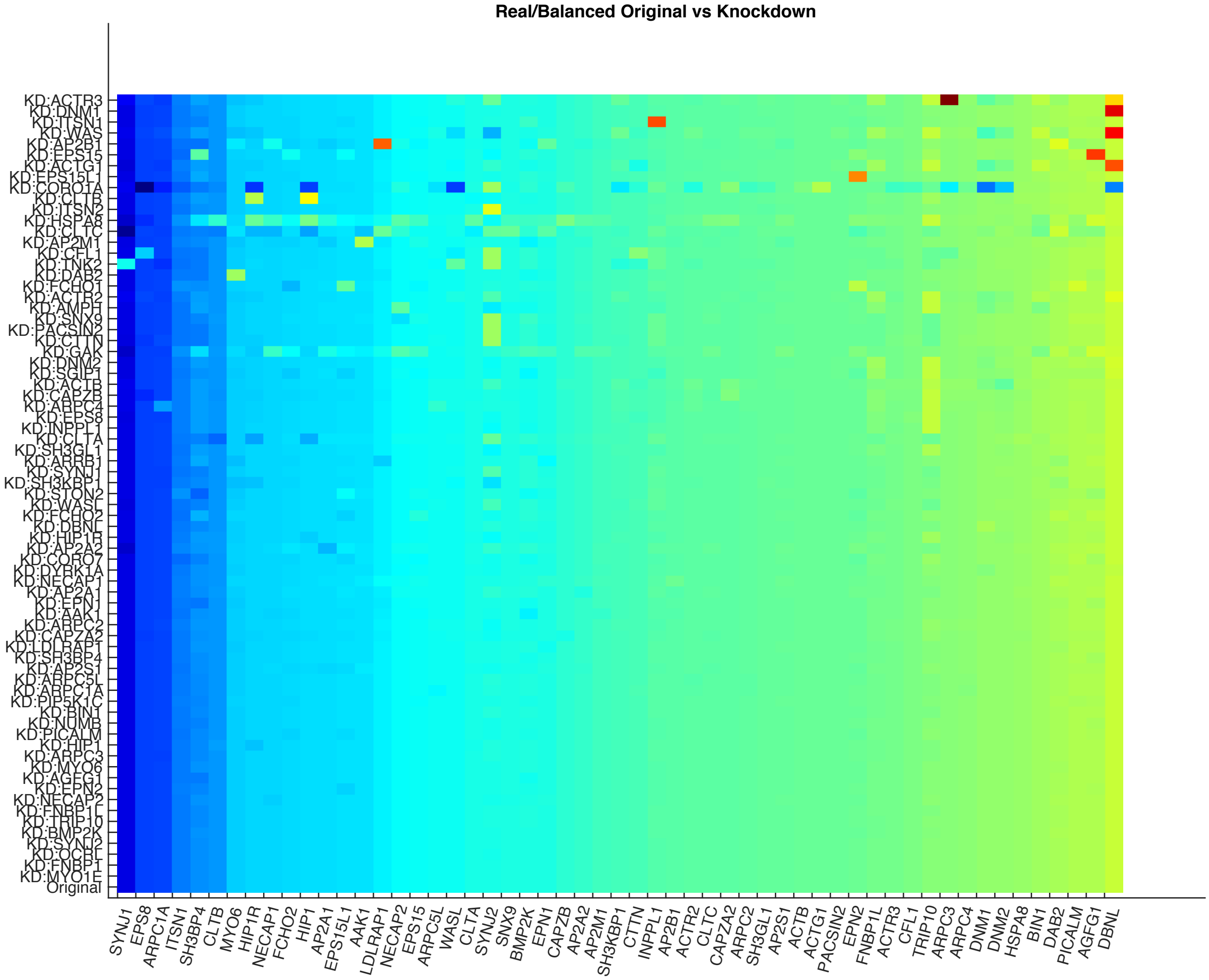


**Fig S14. In fibroblast cells, SB ratios change when nodes are removed or 'knocked-down' (KD) from the network, with most impactful KDs at the top.** The analysis here is for the cytosolic network only. Knock-downs introduce larger perturbations (red spots) relative to the original distribution, in several cases due to proteins with unknown copy numbers.

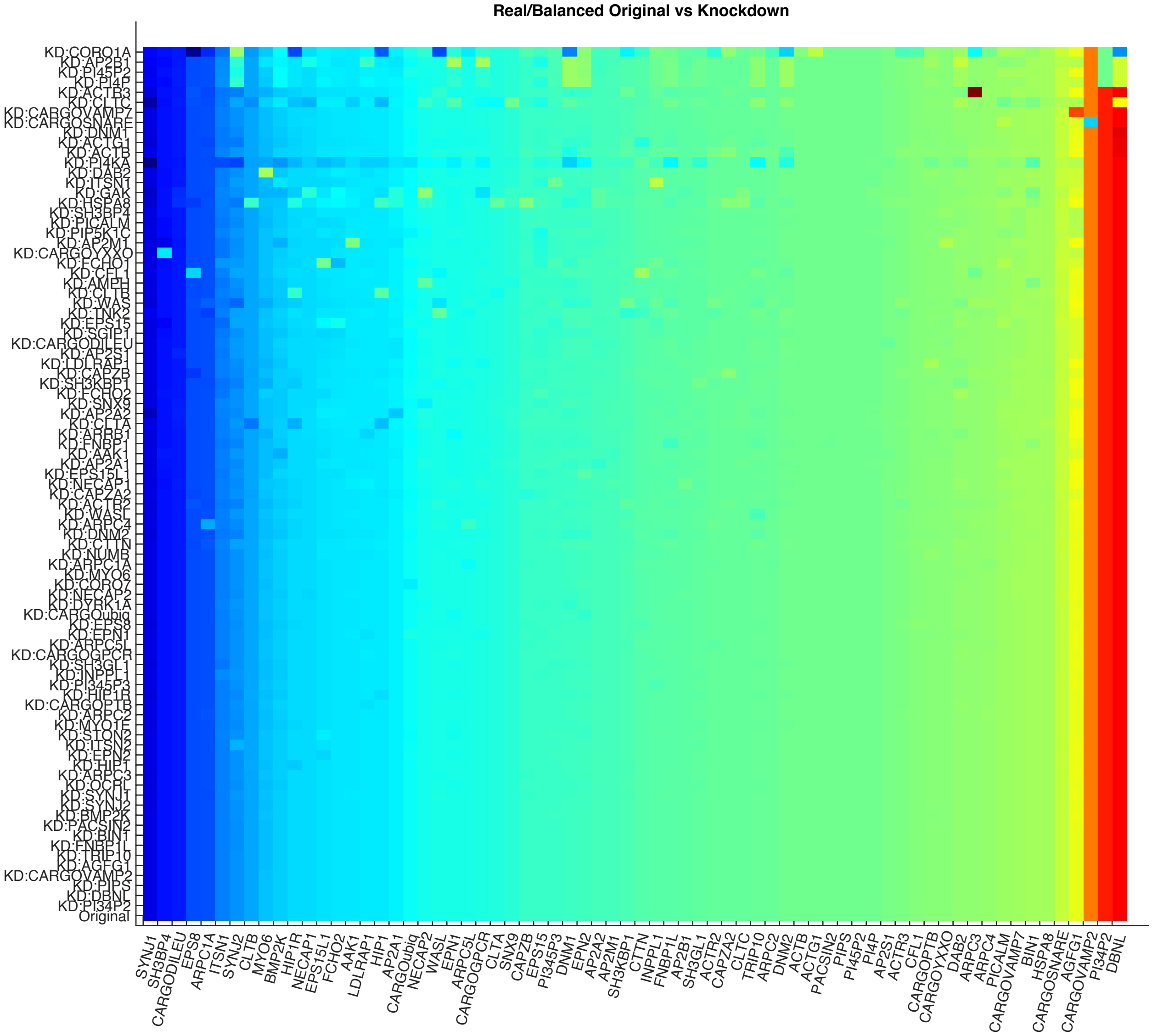


**Fig S15. In fibroblast cells, SB ratios are sensitive to knock-downs.** Here the network includes proteins+cargo+lipids. The lipids stoichiometry for PI/PS was shifted to 1:10, and for AP-2:PIP2 to 1:2. The results are very similar to the 1:1 lipid stoichiometry, except the impact of the AB2B1 removal is higher.

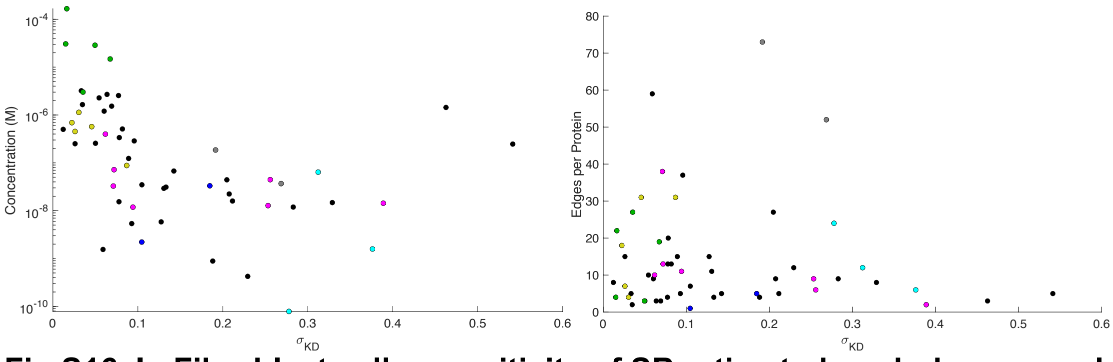


**Fig S16. In Fibroblast cells, sensitivity of SB ratios to knock-down correlates with concentration.** In fibroblast cells, left) the stability or sensitivity relative to concentration has R=-0.45, and with lipids include, R=-0.46. Right) sensitivity correlates only poorly in fibroblast cells with edge number, in part because of the number of proteins with unknown abundance (R=-0.07).

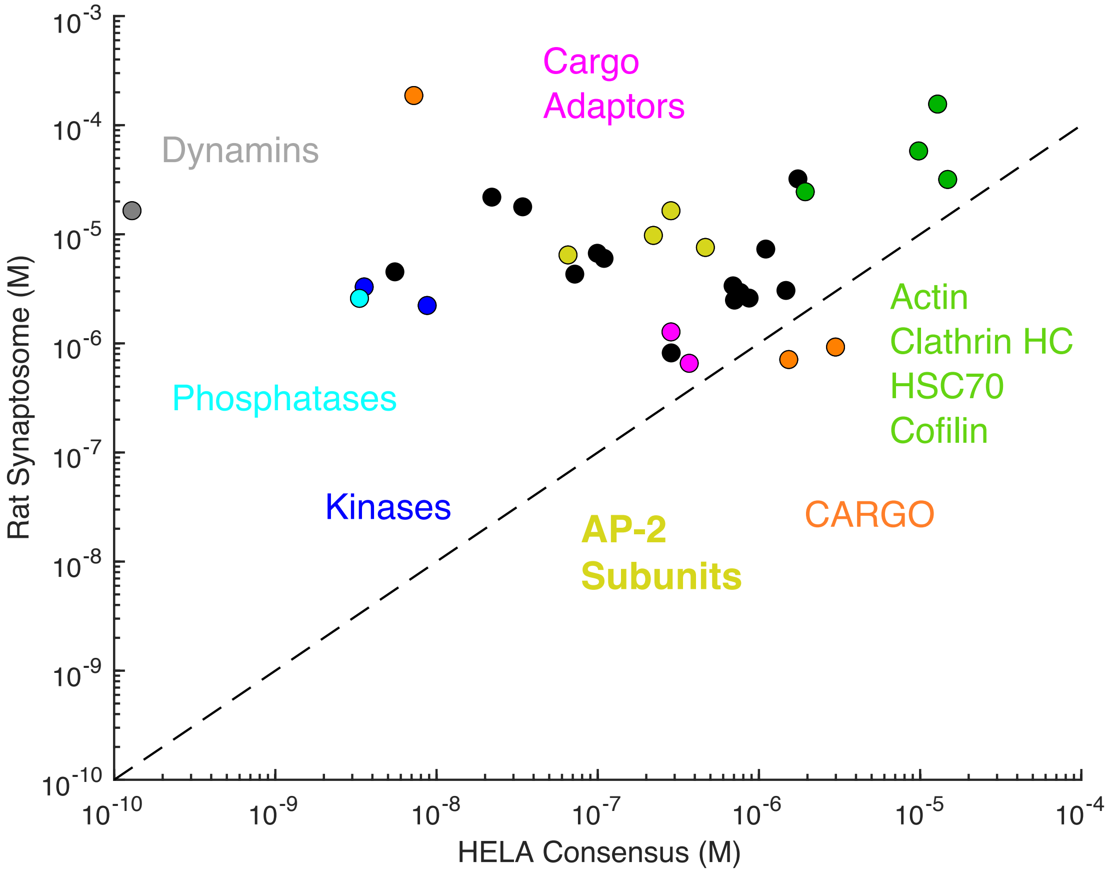


**Figure S17.** Correlation between protein abundances in HeLa cells and Synaptosomes is weak, with R=0.05. This is in large part due to the limited copy number scale of the synaptosome copies.  Only the highly-expressed proteins show similar correlations.  As a result, the many binding partners of AP-2 will greatly outnumber it, relative to the situation in the HELA cells, where AP-2 is relatively abundant compared to many of its partners.

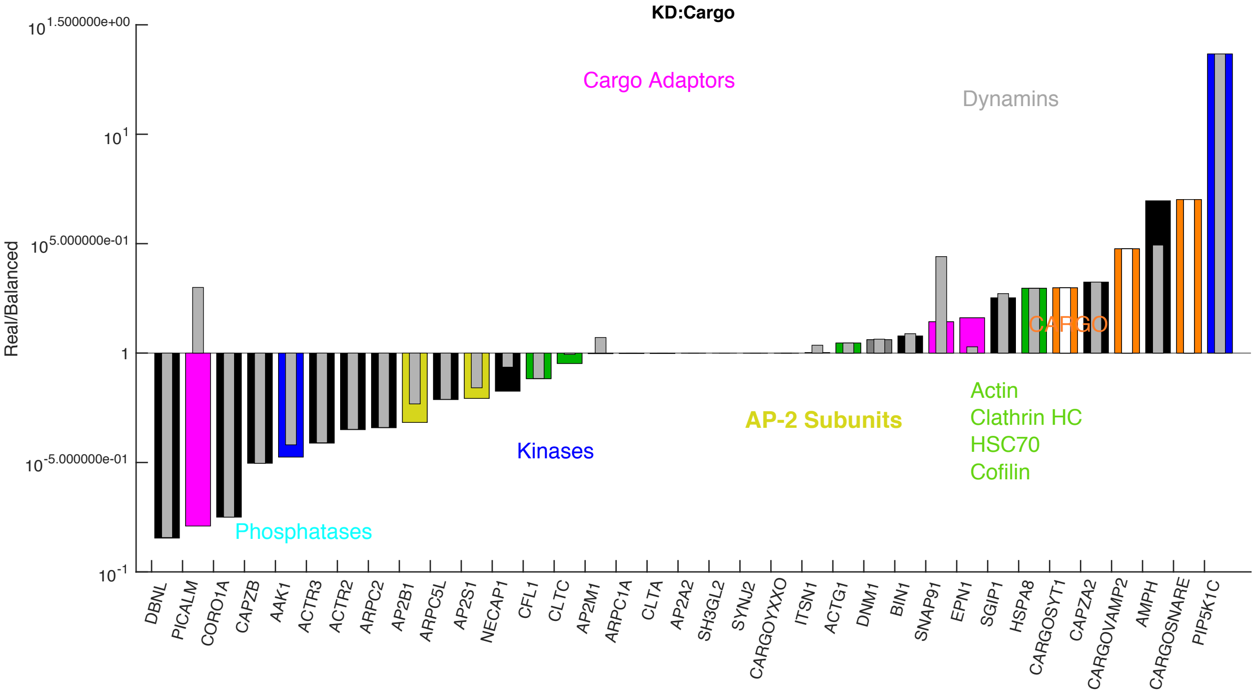


**Figure S18. SB distribution in the synaptosome, network with proteins+cargo.** Many proteins in the synaptosome have unknown abundances, hence the smaller range of SB ratios measureable. Upon KD of the cargo proteins, the SB ratios of the adaptor proteins are shifted (gray bars) especially PICALM. This is the only network where DBNL is sub-stoichiometric. But it is highly unstable in its SB ratio, and with addition of lipids, it returns to being super-stoichiometric.

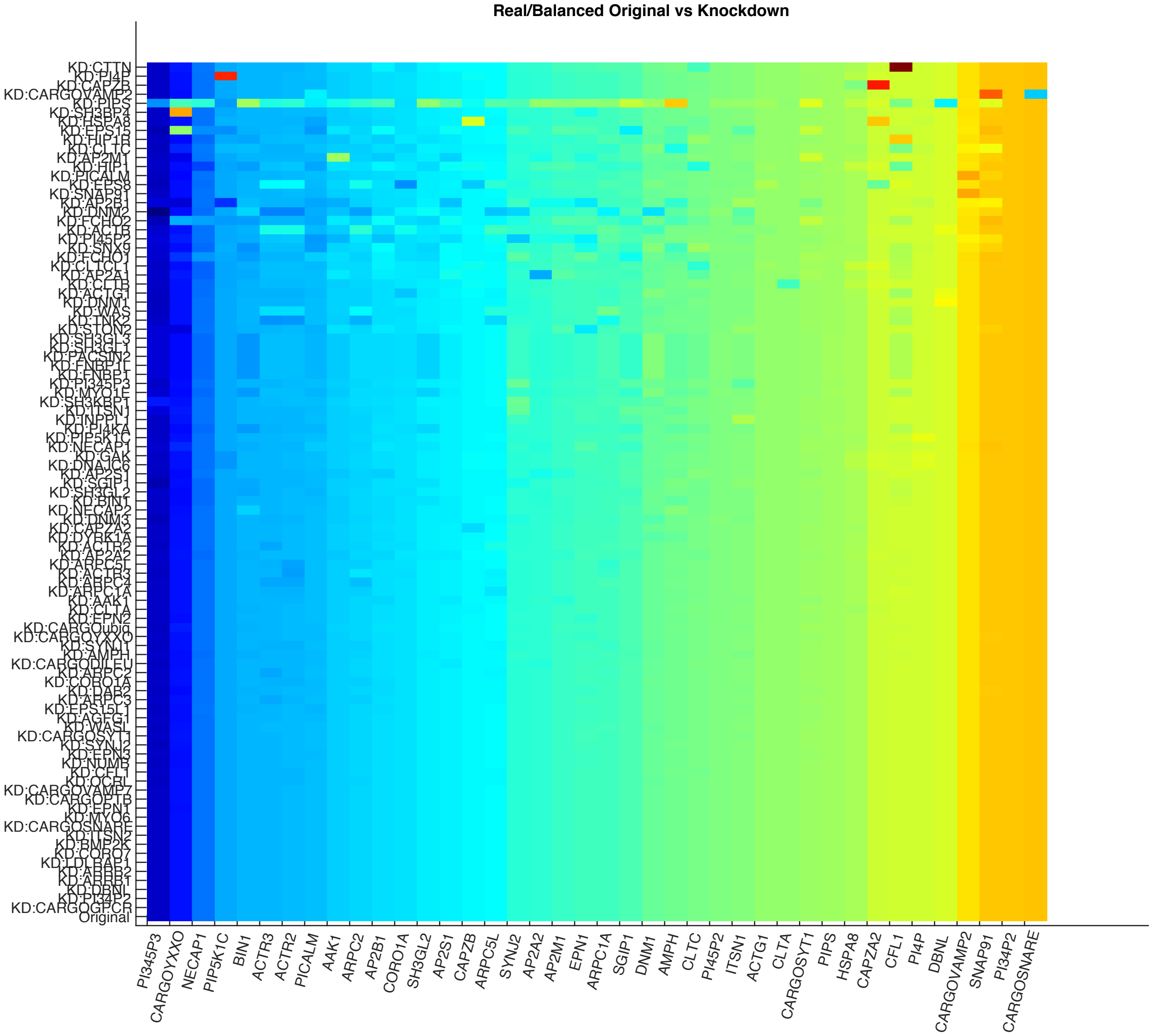


**Figure S19. Matrix of SB ratios in synaptosome, with KD of each protein in the network.** Network with proteins+cargo+lipids. The SNARE cargo SB ratio (far right hand column) is now sensitive to knock-down of the highly expressed VAMP2 cargo, as they both bind to the same adaptor proteins.

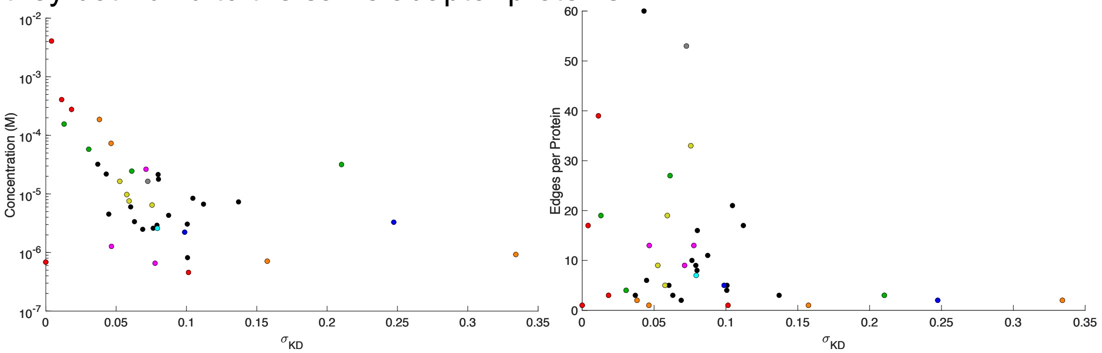


**Figure S20. Sensitivity of stability is anticorrelated with abundance and connectivity in synaptosome.** Left) Synaptosome abundances, with lipids R=-0.63, without, R=-0.43. Right) Edges per protein correlates with R=-0.29, and without lipids, R=-0.24.

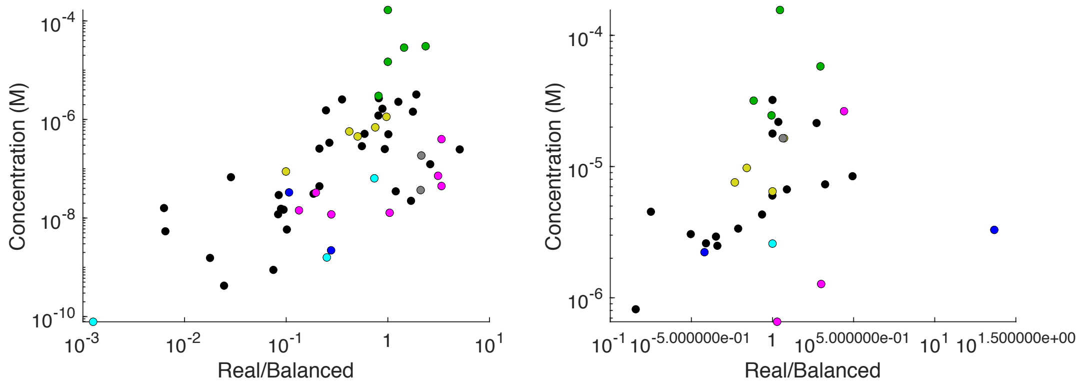


**Figure S21. SB ratios correlate with protein abundance in all cell types.** Left) Fibroblast has R=0.63, with membrane, R=0.56. Right) Synaptosome has R=0.29, with mem, R=0.43.

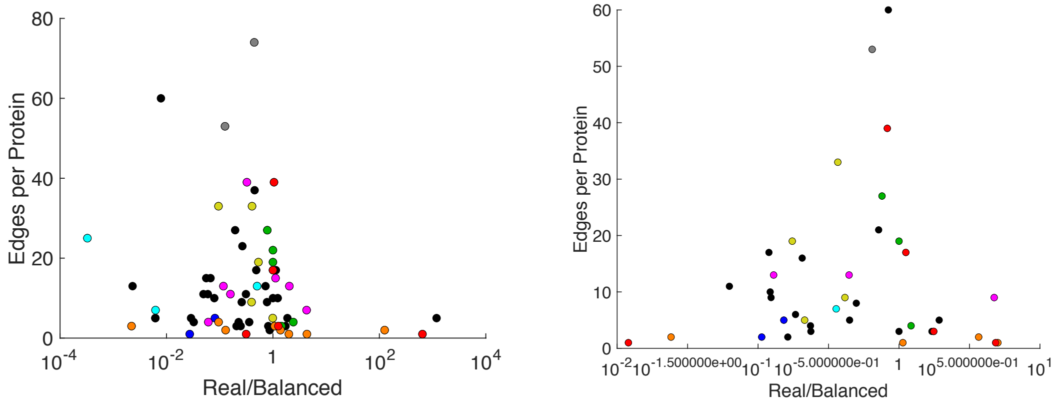


**Figure S22. SB ratios correlate with connectivity, in a volcano plot, across cell types. Concentration and Edge number, by cell type.**

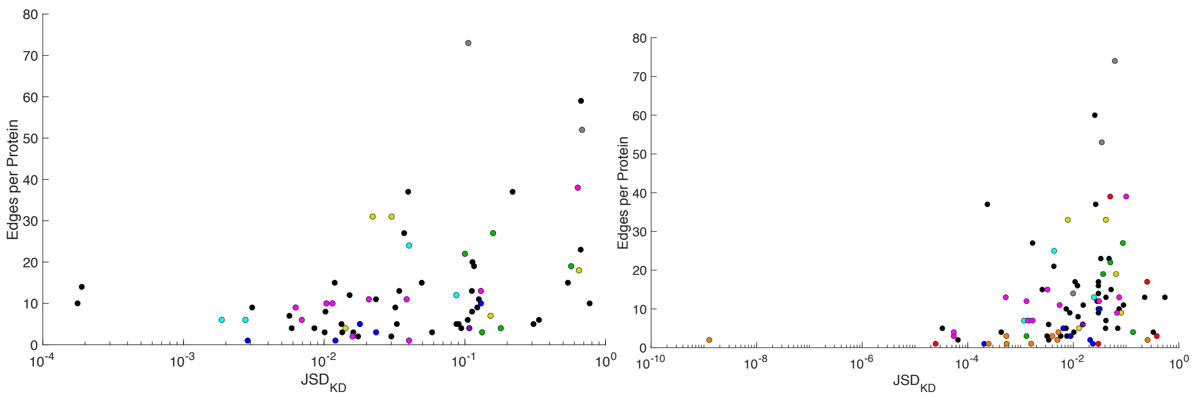


**Figure S23. Disruptiveness of proteins vs connectivity correlates in all cell types.** Left) Fibroblast, R=0.37. With lipids, R=0.21. Right) Synaptosome. R=0.58, without lipids, R=0.29. Positive correlation also exists between JSD and concentrations. For fibroblast, we have R=0.18/0.34 with and without lipids. For synaptosome, we have R=0.28/0.1 with and without lipids.

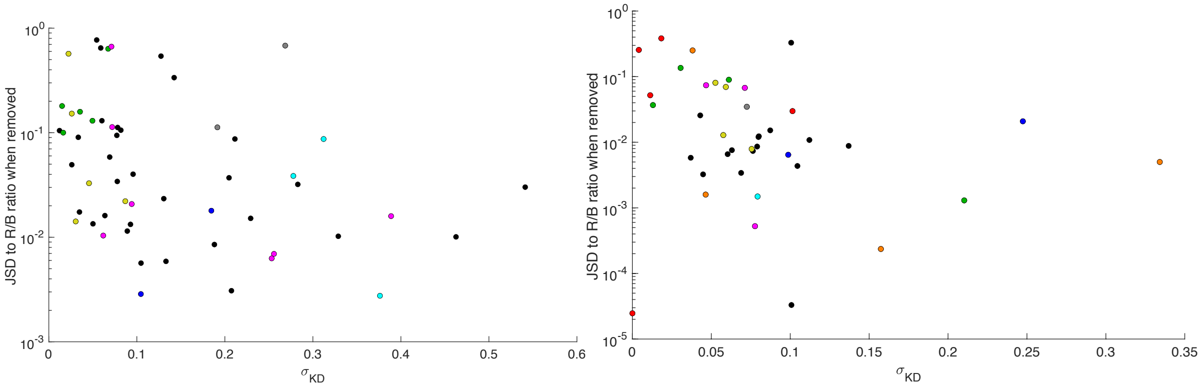


**Fig S24. Proteins that are disruptive to SB distributions are anti-correlated with proteins that have sensitive or unstable SB ratios in all cell types.** Left) Fibroblast, R=-0.35. With lipids, R=-0.57. Right) Synaptosome, R=-0.22, without lipids, R=-0.16.

**TABLES:**

**Tables S1-S3 are in a separate excel file:** SupplementalTablesS1-S3.xlsx

**TABLE S4: HeLa V=1455 $um^3$** **Orange cells have cargo > adaptor**

| **CARGO** | **HeLa Conc (uM) (if cytoplasmic)** | **CARGO Adaptor** | **HeLa Conc (uM)** |
|---|---|---|---|
| SYT1 | 0 | STON2 | 0.0018 |
| VAMP2 | 0.0072 | PICALM+SNAP91 | 0.28+0 |
| GPCR | 0.0599 | ARRB1+ARRB2 | 0.0652+0.0791 |
| Ubiq | 0.1315 | EPN1,2,3+EPS15 | 0.37+0.095+UNK+0.0935 |
| VAMP7 | 0.1744 | AGFG1 | 0.2723 |
| DILEU | 0.2432 | AP2A1+AP2A2+AP2S1 | 0.32+0.065+0.22 |
| PTB | 0.6708 | DAB2+LDLRAP+NUMB | 1.005+0.011+0.045 |
| SNARE | 1.5311 | PICALM | 0.2848 |
| YXXO | 2.9863 | AP2M1+SH3BP4 | 0.28+0.0016 |

**TABLE S5. Fibroblast V=1200 $um^3$** **Orange cells have cargo > adaptor**

| **CARGO** | **Fibroblast Conc (uM) (if cytoplasmic)** | **CARGO Adaptor** | **Fibroblast Conc (uM)** |
|---|---|---|---|
| SYT1 | 0 | STON2 | UNK |
| VAMP2 | 0.0086 | PICALM+SNAP91 | 0.4+0 |
| GPCR | 0.04 | ARRB1+ARRB2 | UNK+0 |
| Ubiq | 0.012 | EPN1,2,3+EPS15 | 0.012+0.013+0+0.033 |
| VAMP7 | 0.04 | AGFG1 | 0.04 |

| | | | |
|---|---|---|---|
| DILEU | 0.005 | AP2A1+AP2A2+AP2S1 | 0.088+0.57+1.33 |
| PTB | 0.39 | DAB2+LDLRAP+NUMB | 0.07+0.014+UNK |
| SNARE | 1.66 | PICALM | 0.4 |
| YXXO | 1.93 | AP2M1+SH3BP4 | 0.45+0.0004 |

**TABLE S6. Synaptosome V=0.235um$^3$ Orange cells have cargo > adaptor**

| CARGO | Synaptosome Conc (uM) (if cytoplasmic) | CARGO Adaptor | Synaptosome Conc (uM) |
|---|---|---|---|
| SYT1 | 73 | STON2 | UNK |
| VAMP2 | 187 | PICALM+SNAP91 | 1.27+26.4 |
| GPCR | UNK | ARRB1+ARRB2 | UNK+0 |
| Ubiq | UNK | EPN1,2,3+EPS15 | 0.65+UNK+UNK+UNK |
| VAMP7 | UNK | AGFG1 | UNK |
| DILEU | UNK | AP2A1+AP2A2+AP2S1 | UNK+6.46+9.76 |
| PTB | UNK | DAB2+LDLRAP+NUMB | UNK+UNK+UNK |
| SNARE | 0.7 | PICALM | 1.27 |
| YXXO | 0.92 | AP2M1+SH3BP4 | 16.4+UNK |
